# Rapid generation of *C. elegans* single copy transgenes combining RMCE and drug selection

**DOI:** 10.1101/2023.03.05.531207

**Authors:** Michael L. Nonet

## Abstract

I outline a streamlined method to create single copy large genomic insert transgenes using Recombination-Mediated Cassette Exchange (RMCE) that relies solely on drug selection yielding a homozygous fluorescent protein (FP) marked transgene in 3 generations (8 days) at high efficiency (>1 insertion per 2 injected P0 animals). Landing sites for this approach are available on four chromosomes in several configurations which yield lines marked in distinct cell types. An array of vectors permit creating transgenes using a variety of selection methods (*HygR, NeoR, PuroR, unc-119(+)*) that yield lines expressing different colored FP tagged lines (*BFP, GFP, mNG*, and *Scarlet*). Although these transgenes retain a plasmid backbone and a selection marker, the inclusion of these sequences typically does not alter the expression of several cell specific promoters tested. However, in certain orientations promoters exhibits crosstalk with adjacent transcription units. In cases where crosstalk is problematic, the *loxP*-flanked fluorescent marker, plasmid backbone and *hygR* gene can be excised by crossing through germline Cre expressing lines also created using this technique. Finally, genetic and molecular reagents designed to facilitate customization of both targeting vectors and landing sites are also described. Together, the rRMCE toolbox provides a platform for developing further innovative uses of RMCE to create complex genetically engineered tools.

## Introduction

Modern biological research is highly dependent on transgenic animals to interrogate cellular functions. They enable both the manipulation of specific gene function in the organism as well as the expression of genetically encoded sensors and subcellular markers (Palmer et al., 2011; Navabpour et al., 2020). In most model systems creating these tools remains an unreliable, time consuming and imprecise process. *C. elegans* is arguably one of the most easily manipulatable metazoan model systems with robust genetics (Jorgensen and Mango, 2002), high throughput RNAi knockdown technology (Conte et al., 2015), as well as efficient genome manipulation techniques including CRISPR and transposon approaches (Nance and Frøkjær-Jensen, 2019). Nevertheless, even in *C. elegans* creation of many types of transgenic animals remains quite time consuming both because of the effort required to create the transgenic animal per se, but also due to the complexities in adapting sensors and markers often designed in other evolutionary distant mouse and fly model systems.

Currently, several approaches are used to create large-DNA single copy insertions in the genome (Nance and Frøkjær-Jensen, 2019). Transposon insertion is a reliable method for creating insertions at random positions in the genome (Frøkjær-Jensen et al., 2014). The transposon approach is relatively efficient, but the position of the insertion cannot be controlled which can be problematic for some types of tool development. In addition, two systems have been developed for creating site specific insertions. Chromosome break repair-based methods were developed first and rely on creating a double stranded break in the genome by either transposon excision (Frokjaer-Jensen et al., 2008) or CRISPR/cas9 cleavage (Kim et al., 2014; Dickinson et al., 2015; Paix et al., 2015), then repairing the break using a template usually provided in the germline via an injection. Recombination based approaches using the FLP/FRT, loxP/Cre and att/phiC31 recombinase systems (Nonet, 2020; Nonet, 2021; Yang et al., 2022) were developed more recently. These systems take advantage of the ability of recombinases to efficiency and accurately recombine sequence containing recombinase sites from a circular donor plasmid into genomic “landing sites” containing compatible recombinase sites. The major advantage of the chromosomal break method is that an insertion can be targeted almost anywhere in the genome. The major disadvantage of the chromosomal break methods is that multiple cellular mechanism can repairs the break, some of which are of low fidelity and introduce errors (Frokjaer-Jensen et al., 2008; Au et al., 2019). The major advantage of the recombinase method is that insertion occurs without invoking a low fidelity mechanism. However, this is at the cost of severe constraints in the position of the insertion.

The current efficiencies of insertion using the CRISPR/cas9-based and the recombinase-based methods are roughly similar. Nonet (2020; 2021) report frequencies between 0.3 to 1.0 of injected animals using recombination and El Mouridi et al. (2022) reports a frequency of 0.25 to 0.75 of injected animals for CRISPR based insertions. In reality, such differences in frequency are largely inconsequential to the time commitment required to develop transgenic lines since the major investments of time in performing an injection are setting up the microscope, preparing the DNA sample (mixing and spinning the sample), loading, mounting, and adjusting the flow of the needle (usually by breaking the tip). Whether one, two or five animals must be injected adds only a small additional time commitment to the process. Thus, further improvements in insertion frequency will provide negligible overt reduction in the time commitment required to perform the injection step of a transgenesis protocol. Although improving insertion frequency remains worthwhile (especially if multiplexed injections are possible), the reality is that developing less complex methodology for introduction of DNA into *C. elegans* is what is required to further reduce the time commitment for this step of the transgenesis pipeline.

Instead, this work is focused on designing vectors and landing sites that reduce the overall time commitment needed to create a transgenic animal by reducing the post-injection manipulations required to identify an insertion, to confirm the structure of an insertion, and to make additional modifications to an insertion. In addition, the pipeline described herein was designed to provide great flexibility in the markers used both to identify an insertion and genetically track an insertion.

The approach uses landing sites that are FP-tagged in a tissue specific manner and express FLP recombinase. DNA sequences of up to 15 kb are inserted into a targeting vector, which when integrated provides drug resistance to select for the insertion, and a change in the FP associated with the tissue specific promoter at the landing site. Insertions can be isolated in 8 days with less than 10 minutes of hands-on time after injection. Furthermore, because several different insertion templates can be coinjected multiple independent isolates of each of three to four distinct transgenes can be routinely isolated from a single microinjection session of 9 animals. The organization of the recombinase sites in the plasmid donor and genomic landing site both prevents re-excision of inserted sequence and prevents extrachromosomal array formation. Consequently, virtually every animal isolated is of the expected molecular structure. The resulting transgenes express a tissue specific FP marker and a selection marker which can be used to track them during genetic manipulations. However, if desired, the tissue specific FP and selection markers can be efficiently excised by crossing the insertions with Cre expressing strains described herein.

## Results

### Overview of rapid RMCE

To create transgenes using the rapid RMCE (rRMCE) method described herein, a targeting plasmid is assembled using standard recombinant DNA methods. Typically, the desired expression module is assembled into the targeting vector using SapI Golden Gate cloning which facilitates coassembly of multiple distinct DNA fragments in an efficient single tube reaction (Figure 1A). In the example outlined in Figure 1, the *lys-8* promoter (*lys-8p*) driving *Scarlet* fluorescent protein (FP) and a *tbb-2 3’* UTR control region was inserted in a *hygR nls-mNeonGreen (nls-mNG*) rRMCE integration vector. The resulting targeting plasmid was then injected into a landing site strain. In the example in Figure 1, a landing site expressing both a nuclear localized cyan-excitable Orange Fluorescent Protein (*nls-cyOFP*) in pharyngeal muscle under the *myo-2* promoter (*myo-2p*) and FLP recombinase in the germline under the control of the *mex-5* promoter (*mex-5p*) was used (Figure 1B, D). Three days after injection, hygromycin was added to the plates, and five days later the plates were screened for homozygous integrants which were identified as viable animals that express a distinct fluorescent marker other than *nls-cyOFP (nls-mNG* in this example). In most cases these animals are the desired final insertion line, which in this example express *Scarlet* under the *lys-8p* robustly in the pharyngeal gland cells and in the intestine in an anterior-posterior gradient (Figure 1B, E). However, if desired, the selection marker (*hygR*), the visual marker (*myo-2p::nls-mNG*) and the plasmid backbone can easily be removed by an outcross to a germline Cre expressing strain requiring two additional generations (Figure 1C, F). Using this approach marked transgenes can be created in as little as 8 days with only 40 minutes of hands-on time (30 minutes for injection, 5 minutes to add the selection agent on day 3 and 5 minutes to clone the homozygous animals on day 8), and unmarked lines can be isolated in 14 days.

**Figure 1.**
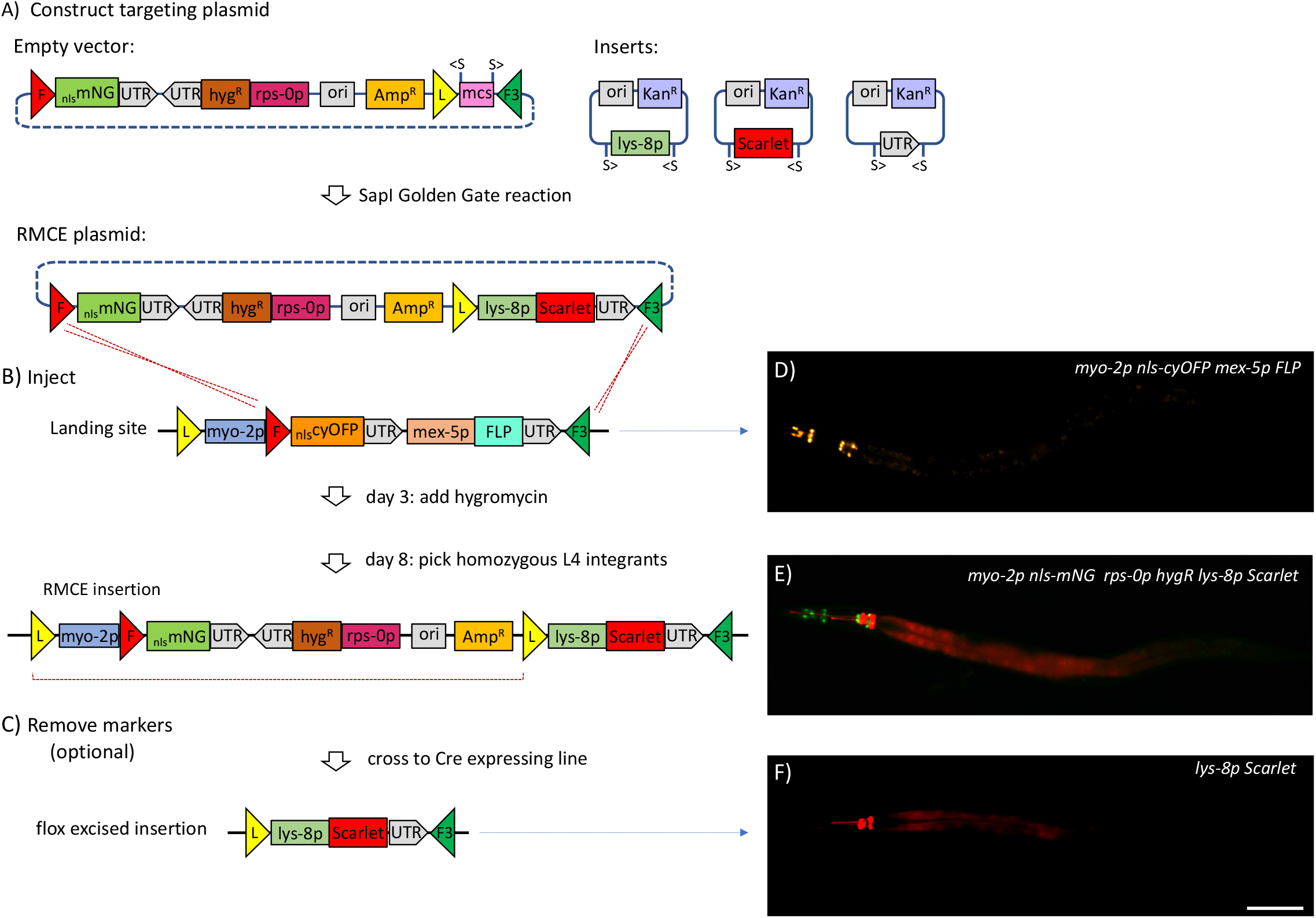
Overview of the rapid RMCE strategy. **A)** A targeting plasmid was constructed using a SapI Golden Gate assembly reaction using a *nls*-mNG targeting vector and clones encoding the *lys-8p*, the fluorescent protein *Scarlet* and the *tbb-2* 3’ UTR. S symbols represent *SapI* restriction endonuclease sites and > symbols indicate the orientation of sites in the plasmids. Colored triangles represent *loxP* (yellow), *FRT* (red) and *FRT3* (green) recombinase sites. **B)** The resulting targeting plasmid was injected into nine adult animals of a landing site strain on Chr II. The landing site strain expresses *nls-cyOFP* in the pharynx and FLP in the germline. The injected animals (3 animals/plate) were incubated at 25°C. On day 3, hygromycin was added to each plate. On day 8, animals with green pharyngeal nuclei homozygous for the insertion were isolated from each plate. **C)** To remove the *myo-2p FP* marker and *hygR* selection cassette, the strain was crossed to a constitutive germline *mex-5p Cre* line. Homozygous animals lacking the floxed *myo-2p nls-mNG hygR* cassette were isolated at high frequency. **D)** Image of an L4-staged animal homozygous for a *myo-2p-nls-cyOFP* marked landing site. **E)** Image of an L4-staged animal homozygous for an insertion isolated as a *hygR* green pharyngeal labeled animal. **F)** Image of an L4-staged animal homozygous for an insertion with the *myo-2p nls-mNG hygR* cassette excised by crossing through a strain expressing germline Cre recombinase. The injection represented in Figure 1B is I435 in Table S2. Strains shown: D) NM5548 *jsSi1726*, E) NM5809 *jsSi1870*, F) NM5881 *jsSi1926*. Scale Bar: 100 μm.

Simplifying the RMCE method for transgene construction and simultaneously providing more flexibility to mark transgenes with a *cis* linked visible marker required extensive modification of the original RMCE integration protocol (Figure S1 and Nonet, 2020). This, in turn, required significant redesign of both the landing sites and the targeting plasmids.

### Creation of Novel Landing sites

Landing sites are generally created by *de novo* introduction of appropriate DNA fragments into the genome using a miniMos, MosSCI or CRISPR approach (e.g. Nonet, 2020; Nonet, 2021). However, an alternate approach was chosen to create these landing sites that relies on modifying existing landing sites using RMCE. The initial rRMCE landing sites were engineered to express *nls-cyOFP* in the pharynx under the control of the *myo-2p* to facilitate both detection and tracking of modifications to the site during the transgenesis pipeline. *cyOFP* was used because it is easily detected in dissecting microscopes and can be readily distinguished from GFP in the 485 nm excitation channel of dissecting microscopes that utilize a long pass emission filter (Chu et al., 2016). In addition, the landing site also expresses FLP under the control of the *mex-5p*, though FLP is not associated with mNG expression in contrast to the previous generation of RMCE landing sites.

*A* two-step process was designed to engineer the landing sites. In the first step, a current landing site was changed from an *FRT/FRT3* landing site to an *FRT10/FRT3* landing site using the strategy outlined in Figure S2. The resulting transgenic line creates a functional *loxP FRT10 FRT3* RMCE intermediate landing site that expresses mNG strongly in the pharynx. A set of intermediate *loxP FRT10 FRT3* landing sites on Chr I, II, IV, V were created with this approach. Subsequently, *myo-2p FRT cyOFP mex-5p FLP* sequences were inserted into a *loxP FRT10 FRT3* vector and RMCE was used to insert these sequences creating the final landing sites. Note that the position of the *mex-5p* FLP transcription unit in these landing sites is distinct from the original single component RMCE landing sites (Figure S1 and Nonet, 2020), and as a result FLP is excised during RMCE rather during the subsequent Cre excision step. This approach was used to create *rRMCE* landing sites on Chr I, II, IV and V.

### Creation of novel Integration vectors

A set of integration vectors for use with the novel landing sites that feature distinct design features from prior RMCE vectors were developed (Figure S3 and Table S1). In the new vectors the selection marker (*hygR, PuroR, or NeoR*) is positioned adjacent to a FP (*mNG, Scarlet, BFP, or GFP*) and separated from a multiple cloning site (MCS) by the pBluescript KS (+) plasmid backbone (Figure 1A and Table S1). This organization was chosen to maximize the distance between the selection marker and the insert sequences. A *loxP* site was included adjacent to the MCS insertion site to permit excision of the selection and marker genes after RMCE. The *FRT* and *FRT3* recombinase sites are positioned such that the entire targeting plasmid except for the 20 bp between these sites is integrated during RMCE.

### Expression characteristics of rRMCE lines

To determine the expression characteristics of insertions derived using the rRMCE approach, a diverse set of promoters of genes of interest (*goi-p*) driving *mNG* or *Scarlet* were inserted into rRMCE plasmids and integrated into landing sites. As illustrated in Figure 2, the insertions express both the *myo-2p FP cis* marker, and the *goi-p FP* insert in the expected cell type-specific pattern. However, unexpected expression was also observed in some lines. The transgenic animals derived using rRMCE contain three distinct transcription units: *myo-2p FP, rps-0p hygR*, and *goi-p FP*. Promoter crosstalk was observed in multiple transgenes created using rRMCE. To characterize this crosstalk more precisely, multiple different insertions with distinct promoters driving fluorescent proteins with distinct emission properties and subcellular localization were created so that the regulatory elements driving the fluorescent reporter could be assessed relatively unambiguously. Several types of interactions were seen. In some cases, the *myo-2p* regulator elements drove expression of both the *nls-FP* just downstream of the promoter itself and in additional also expressed the non-nuclear *FP* downstream of the *goi-p*. Specifically, this was observed with multiple promoters expressing in distinct tissues (Figure 2A, D, F) including the *mec-4p, mex-5p* and the *ehs-1* promoter (*ehs-1p*) suggesting that the *myo-2p* regulatory elements can interact with the basal promoter of other adjacent cell-specific promoters. In another case, the *myo-3* regulatory elements (the *goi-p*) drove expression of the nls-FP associated with the *myo-2p* (Figure 2C). Finally, in other cases, no interaction between promoter was detected (Figure 2B, E) such as with the *cup-4* promoter (*cup-4p*) and the F49H12.4 promoter (*F49H12.4p*). Thus, adjacent transcription units influenced each other in the context of RMCE insertions and these cross-transcription unit influences are dependent on the specific promoters present in the insertion. It appears unlikely that these cross transcriptional unit influences are unique to RMCE insertions, but rather that the design of the insertions made these influences more easily detectable.

**Figure 2.**
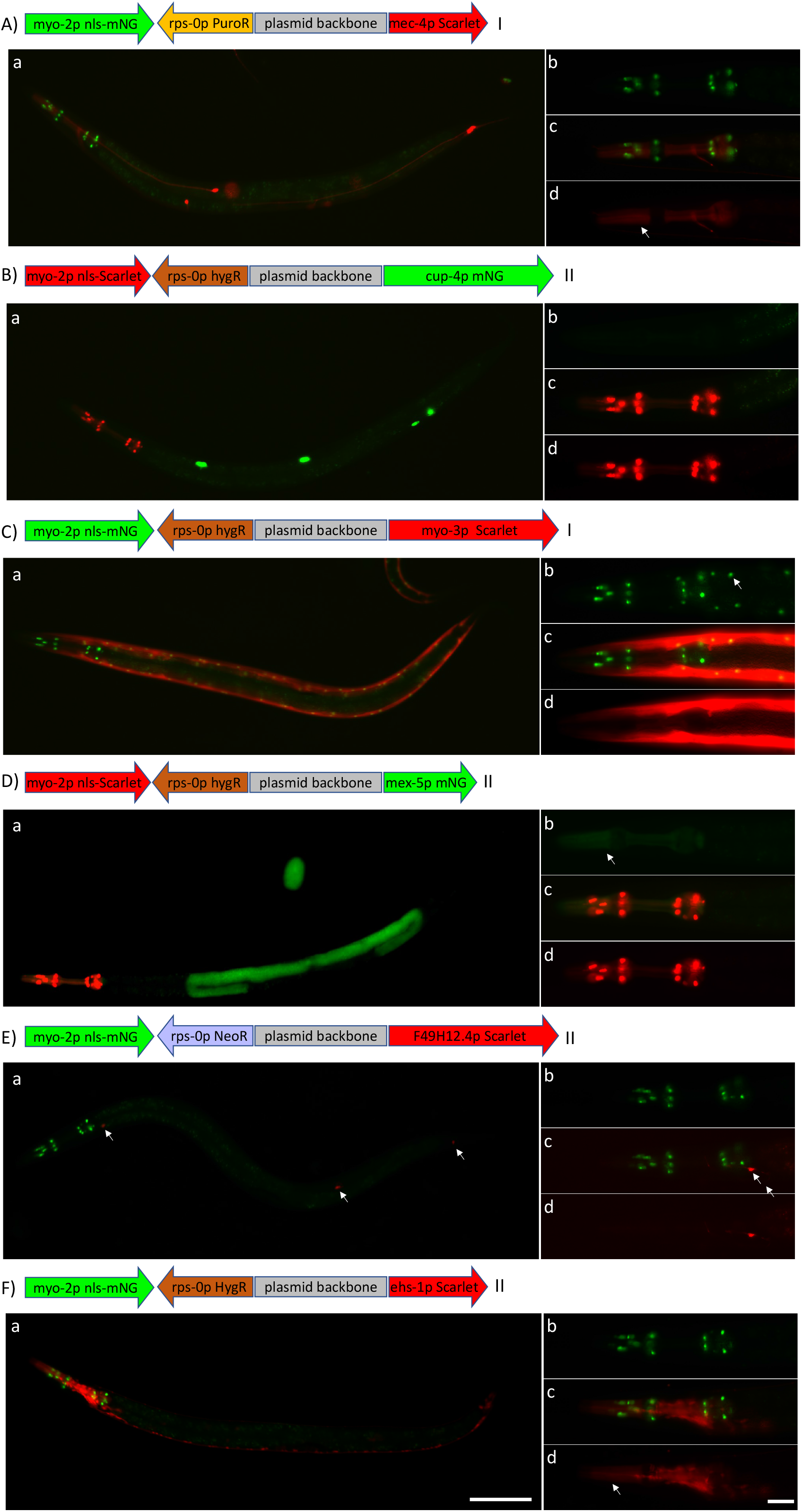
Examples of rapid RMCE insertion transgenes. **A-F)** Images of representative L4-staged rRMCE insertion transgenic animals. Shown on top of each set of images is a diagram of the insertion depicting each transcription unit and the plasmid backbone (to scale) as well as the chromosome location of the insertion. **a)** Widefield view of an animal showing *mNG* (green) and Scarlet (red) fluorescent signals. **b-d)** High magnification view of the pharynx of another representative individual: (b) the *mNG* (d) the Scarlet signal and (c) the merge. **A)** The *mec-4p* driving Scarlet in TRNs, **B)** The *cup-4p* driving mNG in coelomocytes, **C)** The *myo-3p* driving Scarlet in body wall muscle. **D)** The *mex-5p* driving mNG in the germline. An egg is also visible in the image. **E)** The *F49H12.4p* driving Scarlet in the AFD neuron (arrow in head region), the PVD neuron (arrow in mid body), and the PQR neuron (arrow in tail region). **F)** The *ehs-1p* driving Scarlet in a pan-neuronal expression pattern. Note the promoter crosstalk (arrows) seen in Ad, Cb, Db and Fd. Strains shown: A) NM5766 *jsSi1842*, B) NM5770 *jsSi1847*, C) NM5724 *jsSi1827*, D) NM5771 *jsSi1848*, E) NM5822 *jsSi1874*, F) NM5903 *jsSi1940*. Scale Bars: 100 μm (whole animal images) and 25 μm (pharyngeal images).

### Influences of spacing, orientation and position on promoter crosstalk

To further characterize promoter crosstalk interactions, additional integration plasmids were created to examine the effects of the orientation of the transcription units in the insertion on crosstalk. RMCE integration plasmids were constructed with the *rps-0* promoter (*rps-0p) hygR* selection marker in reverse orientation, and versions of both hygR orientations with the MCS re-oriented. Integration plasmids containing the germline *mex-5p* driving mNG and a *mec-4p* driving *Scarlet* were created in all 4 orientations and inserted on Chr II. When all three transcription units were oriented in the same direction strong cross-promoter interactions were observed with *rps-0p* (and probably also *myo-2p*) expressing the FPs associated with the *mex-5p* and *mec-4p* as evidenced by the non-nuclear FP signal broadly visible in most somatic tissues and robustly observed in the pharynx (Figure 3A, E). Crosstalk was still present, although greatly reduced, when the *rps-0 HygR* transcription unit was flipped (Figure 3B, F) and was virtually (*mex-5p*) or totally (*mec-4p*) eliminated when the gene of interest transcription unit was flipped (Figure 3C, D, G, H). This cross was not specific to the Chr II insertion, occurring equally robustly for insertions on Chr I, IV and V (Figure S4).

**Figure 3.**
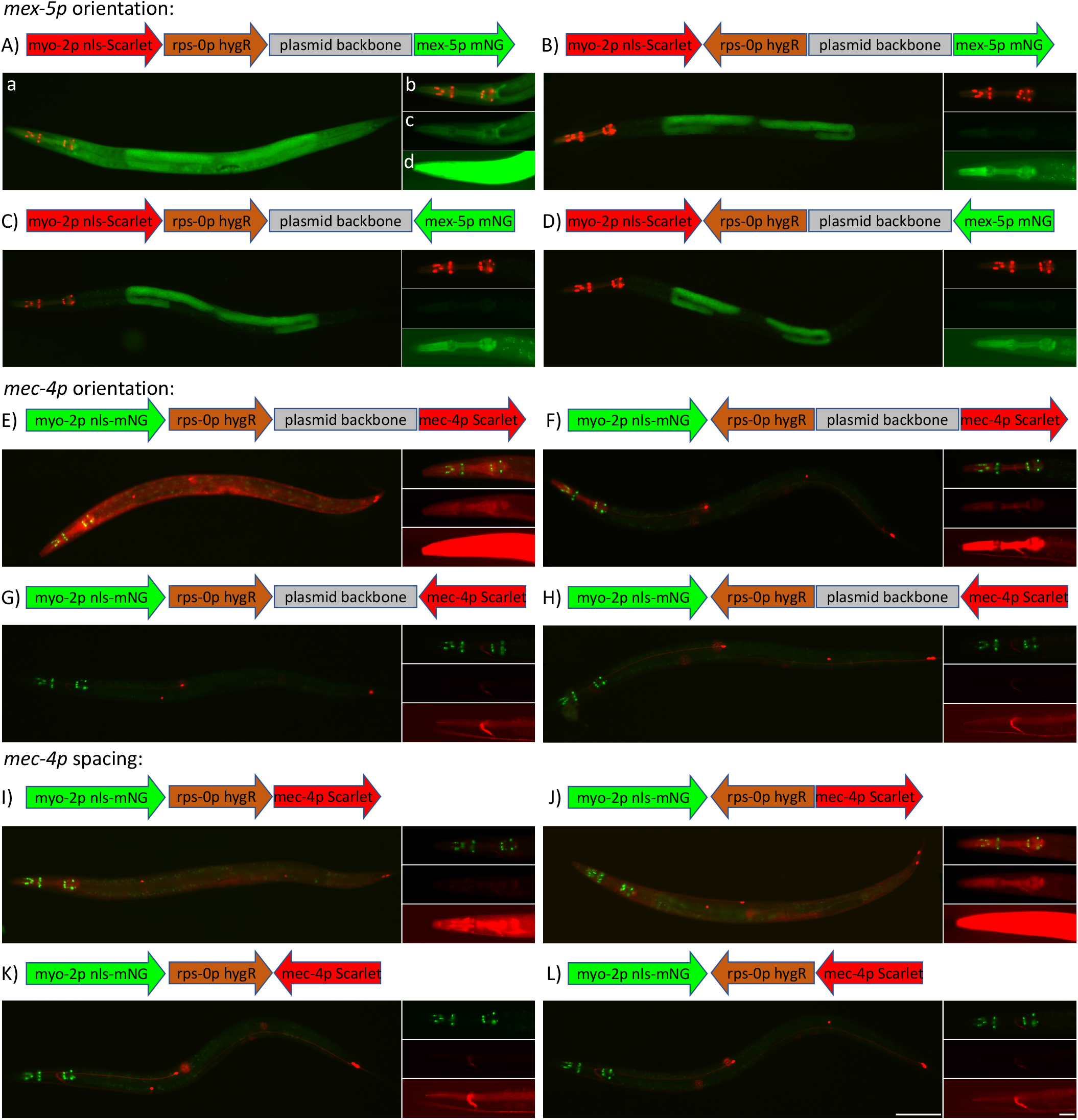
Influences of transcription orientation and spacing on the specificity of expression in rRMCE transgenes. **A-L)** Images of representative L4-staged rRMCE insertion transgenic animals. Shown on top of each set of images is a diagram of the insertion depicting each transcription unit and the plasmid backbone (to scale). **a)** Widefield view of an L4 animal showing mNG (green) and Scarlet (red) fluorescent signals. **b-d)** High magnification view of the pharynx of another representative individual: (b) a merge of the mNG and Scarlet signals, (c) the reporter signal (d) and a long exposure of the reporter signal. Note the crosstalk seen in Ad, Bd, Ed, Fd, Id, and Jd. Subpanels are only labelled in A. **A-D)** Effects of orientation on expression of a *mex-5p* mNG reporter integrated on Chr II in a *myo-2p* marked landing site. **E-H)** Effects of orientation on the expression of a *mec-4p* Scarlet reporter integrated on Chr II in a *myo-2p* marked landing site. **I-L)** Effects of spacing between transcription units on the expression of a *mec-4p* Scarlet reporter integrated on Chr II in a *myo-2p* marked landing site. Strains shown: A) NM5870 *jsSi1910*, B) NM5771 *jsSi1848*, C) NM5853 *jsSi1914*, D) NM5768 *jsSi1845*, E) NM5824 *jsSi1878*, F) NM6018 *jsSi2025*, G) NM5804 *jsSi1862*, H) NM5864 *jsSi1917*, I) NM5873 *jsSi1922*, J) NM5871 *jsSi1920*, K) NM5908 *jsSi1955*, L) NM5910 *jsSi1953*. Scale Bars: 100 μm (whole animal images) and 25 μm (pharyngeal image).

To assess how distance is influencing crosstalk, the position of the plasmid backbone was moved in the RMCE integration vectors from between the *hygR* gene and the insert to between the FRT and FRT3 sites. In this position, the plasmid backbone sequences are not introduced into the genome during integration. *mec-4p Scarlet* was inserted into the four orientation vectors and integrated on Chr II. In the resulting transgenes, loss of the 2.5 kb backbone spacer increased crosstalk when *hygR* and *mec-4p* FP were transcribed in divergent orientations (Figure 3I, J). Loss of the spacer had little effect on crosstalk when *mec-4p FP* was oriented towards the *hygR* gene (Figure 3K, L). In summary, both the relative position and orientation of the transcription units influence expression in rRMCE transgenes.

While the *myo-2p* is very useful for marking transgenes, the robust influence of the *myo-2p* on the expression of adjacent genes in some single copy insertions suggests that it might not be the ideal promoter to mark transgenic insertions in some situations. For example, even low-level promoter crosstalk could potentially influence the interpretation of a transgenic approach to test the cell-type specificity of gene function. Several approaches were used to mitigate crosstalk including utilizing different promoters for the *cis marker*, creating transgenes that are unmarked, and creating Cre reagents to convert marked insertions into unmarked insertions. To develop these alternatives tools novel rRMCE landing sites were required.

### Additional cell type-specific landing sites

A set of intermediate *loxP FRT10 FRT3* landing sites on chr I, II, IV and V that express both *myo-2p FRT10 mNG* and *mex-5p FLP D5* were used to create new cell type specific landing sites (Figure 4A). In addition, a *loxP FRT10 FRT* RMCE vector containing a *mex-5p FLP* transcription unit was constructed to simplify creating novel targeting plasmids (Figure 4B). These reagents can be used to create a landing site regulated by any promoter of interest using a single cloning step followed by a single RMCE integration (Figure 4B, C).

**Figure 4.**
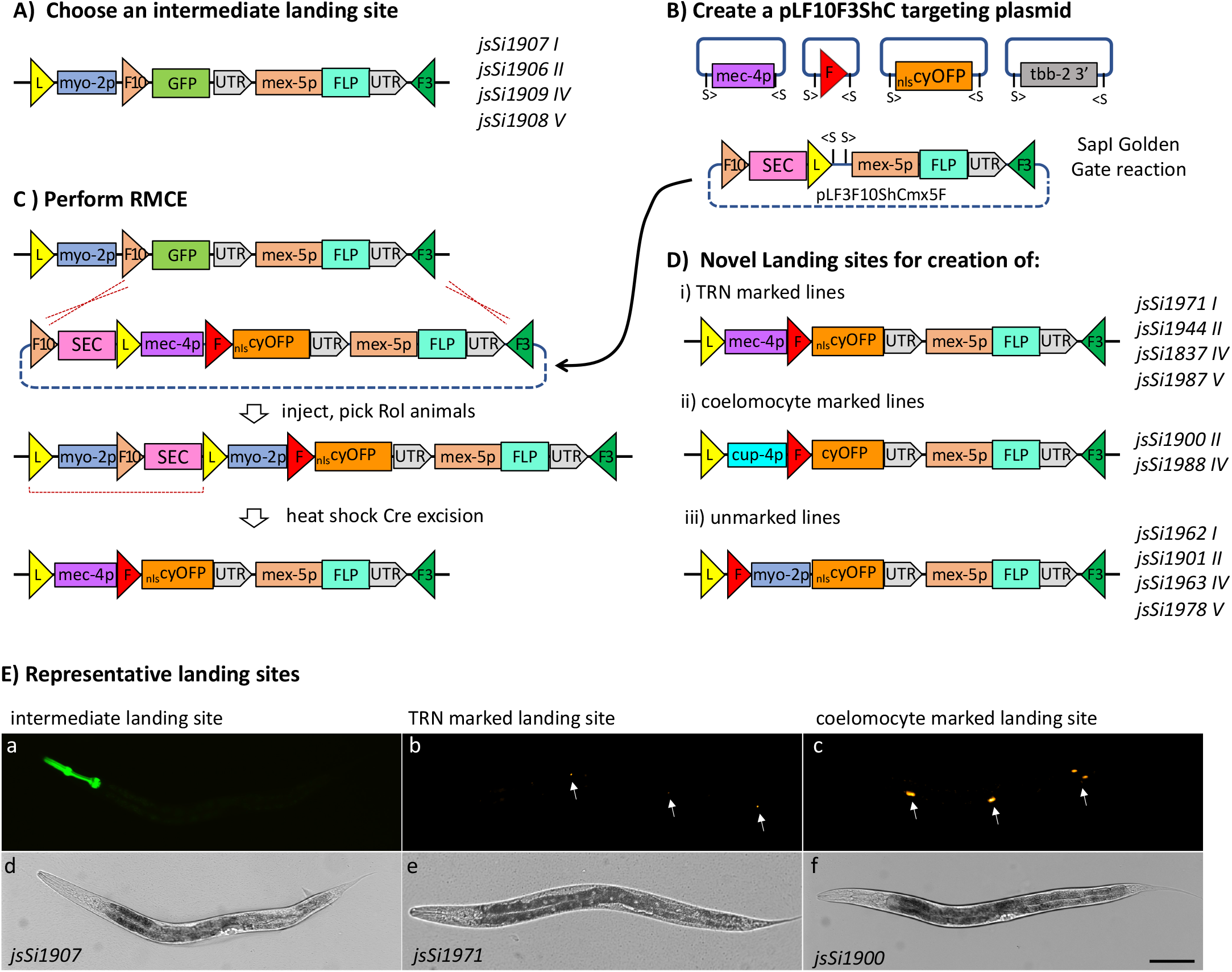
Methodology used to create custom landing sites. **A)** Diagram of the intermediate landing sites created on each of four chromosomes to simplify development of novel rapid RMCE landing sites. These landing sites were created using an RMCE strategy as outlined in Figure S2. The allele designation and chromosome of each insertion is listed on the right. **B)** Diagram of the methodology used to construct the targeting plasmid to create a TRN neuron expressing landing site using a *SapI* Golden Gate cloning approach. The *kanR* plasmids containing the *mec-4p*, an *FRT* site, *nls-cyOFP* and the *tbb-2* 3’ UTR were co-assembled into an *AmpR* rRMCE vector containing a *mex-5p FLP* expression cassette. S symbols represent SapI sites and > symbols indicate the orientation of sites in the plasmids. **C)** Diagram of the RMCE procedure used to integrate the landing site targeting plasmid into the genome to create the novel landing site. A *sqt-1* Rol screening strategy was used to identify insertions as described in Nonet, 2020. Molecular components for building landing sites at new genomic locations are also available (Figure S7). **D)** Diagram depicting the structure of the customized landing sites currently available. The allele designation and chromosome of each insertion is listed on the right. Note that *jsSi1837, jsSi1900* and *jsSi1901* were created using intermediate landing sites similar, but not identical, to those depicted in panel A). See Table S3 for details. **E)** Images of representative landing sites. **Top row)** Widefield epi-fluorescence images of L4-staged animals. Arrows in b) point to ALMR, PVM and PLMR. Arrows in c) point to the three sets of coelomocytes. **Bottom row)** brightfield images of the same animals. Scale Bar: 100 μm. Diagrams of plasmids and chromosomal insertions are not to scale. Colored triangles represent *loxP* (yellow), *FRT* (red), *FRT3* (green) and *FRT10* (light brown) recombinase sites. SEC represents a <u>S</u>elf-Excision <u>C</u>assette consisting of the *hsp-16* promoter driving *Cre*, the *sqt-1* promoter driving *sqt-1(e1350*), and a promoter-less *hygR* gene. Strains shown: NM5861 *jsSi1907*, NM5879 *jsSi1900 and NM5937 jsSi1971*.

The *myo-2p* has long been used as a transformation marker because it is both strong and highly specific. However, for work in the pharynx or the nerve ring, strong signal from pharyngeal nuclei can interfere with imaging a process of interest. Two other less strong, yet specific, promoters were chosen to create the additional cell type-specific landing sites with the idea that 1) these would provide suitable alternatives for experiments where *myo-2p* signals are problematic, and 2) the hope that these promoters might exhibit less crosstalk. *mec-4p* was chosen because it is well characterized and often utilized in my studies. *cup-4p* was chosen because it is relatively strong, specific and did not show crosstalk interactions with the *myo-2p* (Figure 2B). *mec-4p* landing sites at four chromosomal positions and *cup-4p* landing sites at two chromosomal positions were created (Figure 4 D, E). Integration plasmids expressing *mex-5p mNG, mec-4p mNG* (Figure 5 and Figure S5A, B) and other promoters (Figure S6) were introduced into these alternative landing sites using rRMCE and examined to characterize expression and crosstalk. Both *mec-4p* and *cup-4p* landing site behaved analogously to *myo-2p* landing sites, exhibiting high crosstalk when all 3 transcription units were oriented similarly (Figure 5A, C), less crosstalk when the *hygR* was oriented in reverse (Figure 5 E, G and Figure S6 G, H) and little, if any, crosstalk when the *hygR* and reporter transcription units were both in reverse orientation (Figure 5 B, D, F, H). Other promoters behaved similarly (Figure S6 C-F). These landing sites should provide a reasonable alternative approach for creating marked lines, though the expression levels in TRNs and coelomocytes are lower than that of *myo-2p* based lines and thus more difficult to detect in a dissecting microscope. The use of integration vectors (Table 1) that express a cytosolic rather than nuclear localized FP yield significantly higher signals and mitigate this drawback.

**Figure 5.**
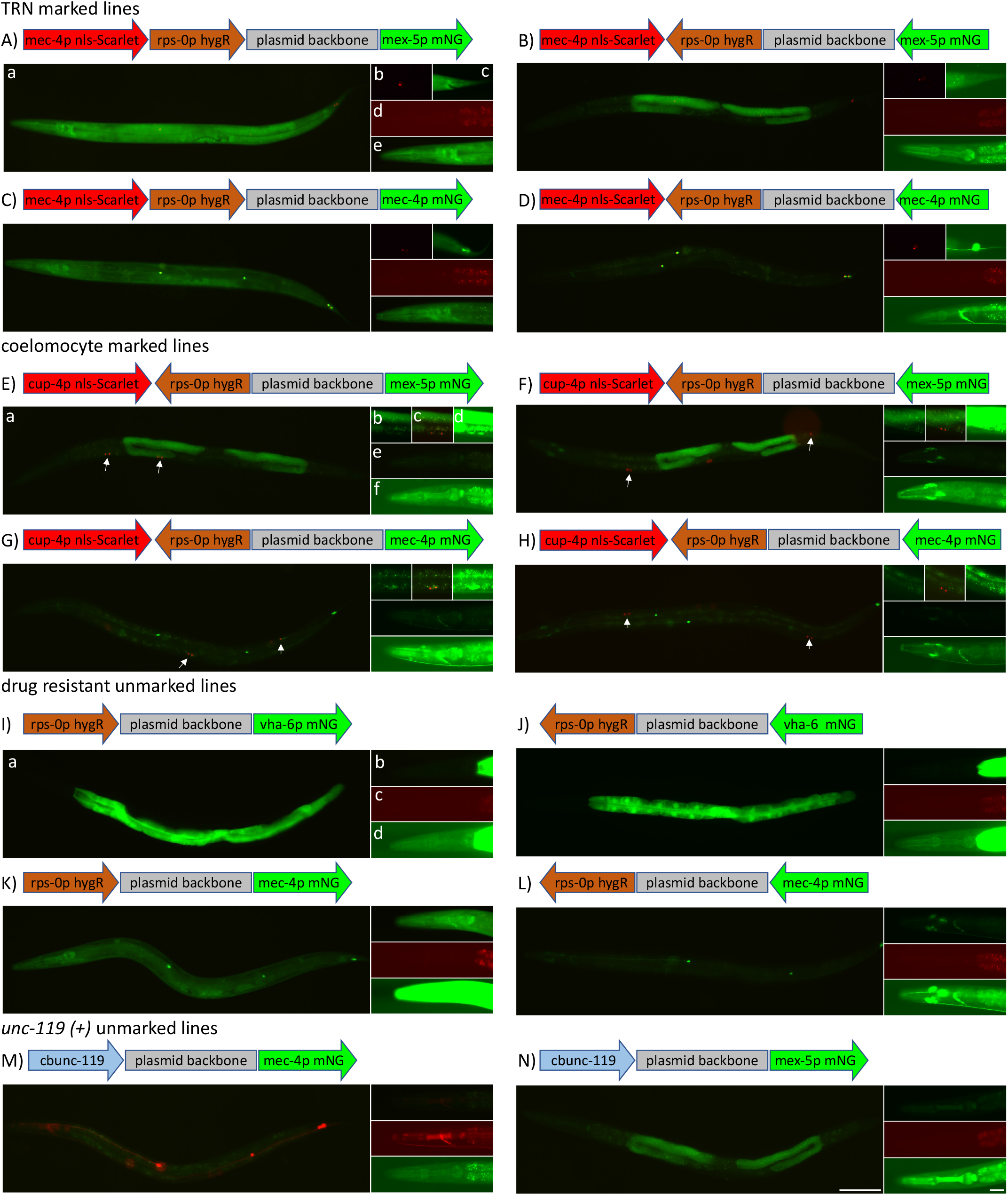
Representative TRN, coelomocyte, unmarked and unc-119(+) rRMCE lines. **A-M)** Images of representative L4-staged rRMCE insertion transgenic animals. Shown on top of each set of images is a diagram of the insertion depicting each transcription unit and the plasmid backbone (to scale). **a**) Widefield view of an L4 animal showing mNG (green) and Scarlet (red) fluorescent signals. **A-D)** TRN marked *mex-5p* and *mec-4p* lines on Chr I. **b-c)** High magnification view of the tail of another representative individual: (b) the Scarlet signal (c) the mNG green signal (a long exposures in B and D). **d-e)** High magnification view of the pharynx of another representative individual: (d) a long exposure of the Scarlet signal (e) the mNG green signal (a long exposures in B and D). **E-H)** Coelomocyte marked *mex-5p* and *mec-4p* lines on chr IV. **b-d)** High magnification images of coelomocytes in the mid-body of a representative L4 animal: (b) the mNG signal, (c) a merge, and (d) a long exposure of the mNG signal. **e-f)** High magnification view of the pharynx of another representative individual: (e) a merge and (f) a long exposure of the mNG signal. **I-L)** hygR resistant unmarked *vha-6p* and *mec-4p* lines on chr IV. **b-d)** High magnification view of the pharynx of another representative individual: (b) a merge and (c) a long exposure of the Scarlet signal, (d) a long exposure of the mNG signal. **M-N)** *unc-119(+*) unmarked *mec-4p* and *mex-5p* lines on chr V. **b-d)** High magnification view of the pharynx of another representative individual: (b) a merge and (c) a long exposure of the Scarlet signal, (d) a long exposure of the mNG signal. Note the unexpected GFP + cells adjacent to the anterior region of the pharynx that are visible Chr IV insertions in F, H, and J. Strains shown: A) NM6123 *jsSi2141* B) NM6149 *jsSi2155*, C) NM6095 *jsSi2124*, D) NM6082 *jsSi2082*, E) NM6052 *jsSi2062*, F) NM6116 *jsSi2143*, G) NM6046 *jsSi2061*, H) NM6146 *jsSi2144*, I) NM6140 *jsSi2138*, J) NM6142 *jsSi2137*, K) NM6131 *jsSi2139*, L) NM6106 *jsSi2129*, M) NM6009 *jsSi2022* and N) NM6008 *jsSi2020*. Scale Bars: 100 μm (whole animal images) and 25 μm (pharyngeal image).

### Creating unmarked lines

Reagents to create lines that are not marked by a *cis*-linked promoter FP fusion were also developed. To make rRMCE amenable to creating this type of line, the *FRT* site in the landing site targeting plasmid was moved to a position 5’ of *myo-2p* and landing sites with this structure were created on four chromosomes (Figure 4D). In this position the RMCE insertion replaces the *myo-2p* creating a promoter-less insertion. Integration plasmids that lack a fluorescent marker 3’ of the *FRT* site were also created as well as a vector that replaces *hygR* with *C. briggsae unc-119* (Table S1). A set of promoter FP combinations were inserted into these vectors and integrated into the new landing sites (Figure S5 C, D). These insertions generally behave similarly to other landing sites with significant promoter crosstalk being observed when the *hygR* and reporter transcription units are similarly oriented (Figure 5K & Figure S6I), while showing little crosstalk when hygR orientation was reversed (Figure 5L & Figure S6J). Interestingly, the *vha-6* promoter (*vha-6p*) and *myo-3p* appear resistant to *hygR* crosstalk (Figure 5 I, J & Figure S6 K, L). The *unc-119* lines were relatively specific though a low level of pharyngeal signal was observed (Figure 5 M, N). These *FRT myo-2p* landing sites provide a suitable alternative for creating unmarked transgenic lines when a *cis*-marker reporter is not desired or potentially problematic.

### Cre-excision of marked lines

In case where even minor promoter crosstalk is of serious concern in interpretation of an experiment, a marker-less transgene may provide more certainly of the specificity of expression of an insertion. An additional set of vectors that lack the *loxP* site between the plasmid backbone and the insertion site were created and used to integrate *mex-5p nls-Cre* constructs into landing sites on all four chromosomes (Table S1 and Figure 6A, B). These insertions only have a single *loxP* site preventing *Cre* from excising the marker and *hygR* transcription units which would circumvent the selection method for rRMCE. The lines permit excision of the FP marker, the *hygR* selection cassette and the plasmid backbone from standard rRMCE insertions (those that contain two *loxP* sites) via a single cross. Excision of the *myo-2p FP hygR* backbone region occurred in over 95% of chromosomes derived from a *trans* heterozygote carrying one copy of a *mex-5p* Cre line and one copy of an insertion being excised (Figure 6C).

**Figure 6.**
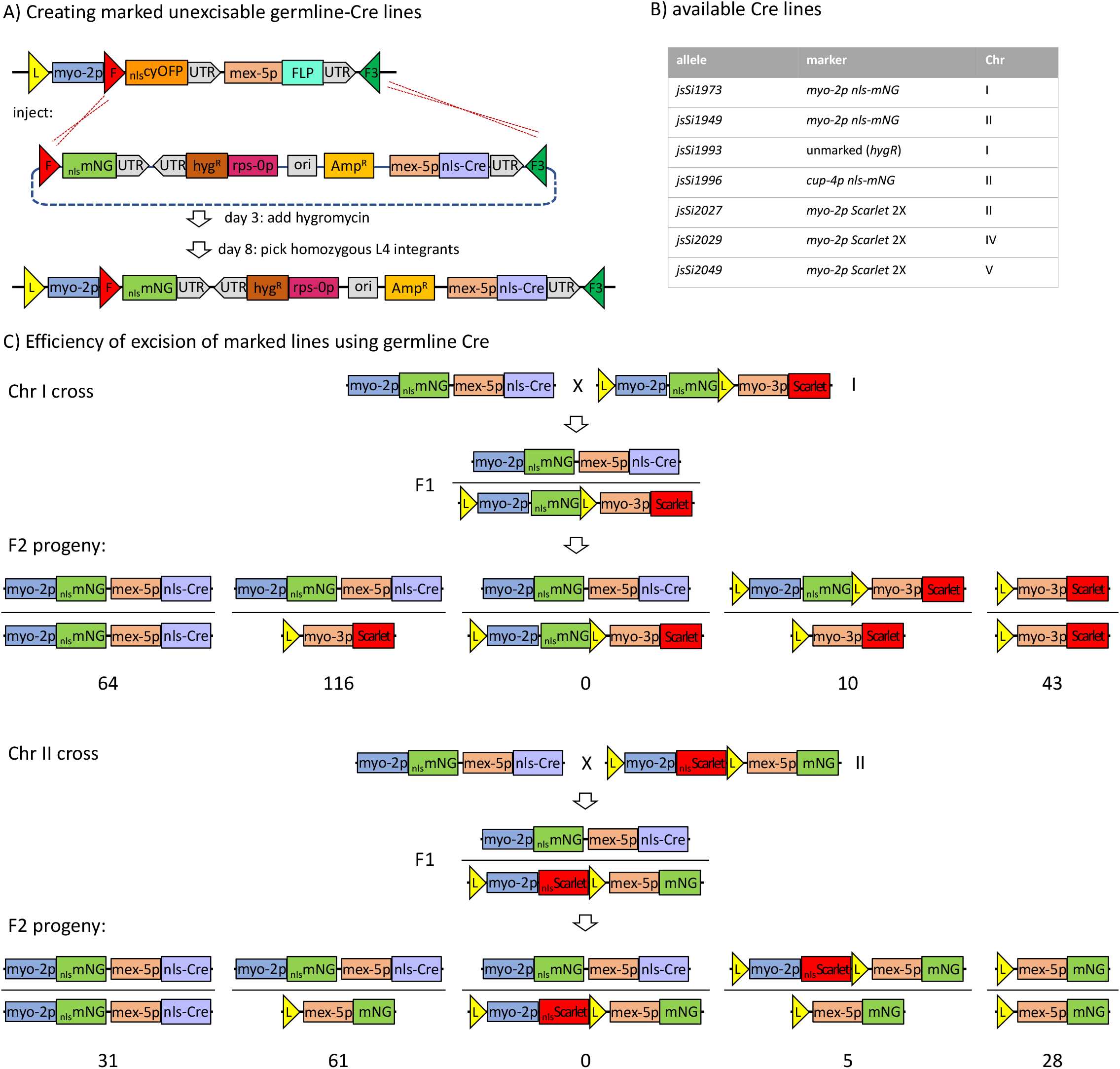
Creating unmarked insertions using Cre excision. **A)** Outline of the strategy used to create germline Cre expressing marked lines. In the example shown, the *mex-5p* driving *nls-Cre* was inserted into the pHygG8 rRMCE vector which lacks the *loxP* site required for excision of the marker and hygR selection cassette. This results in an insertion that cannot be excised by the germline expression of Cre from the transgene. **B)** Several germline Cre lines were created at different landings sites expressing various florescent proteins in several cell-types. The variety permits simple tracking of the Cre line regardless of the markers expressed in the line being excised. See Table S4 for list of strains. **C)** The efficiency of Cre-mediated excision of the marker is high. Two crosses were performed to quantify the efficiency of excision. A simplified schematic of the structure of the chromosomes in the crosses and the number of animals obtained of each genotype are shown. The genotypes of the parent/excised chromosome *trans* heterozygote animals were confirmed by scoring progeny of each of the ten individuals in cross 1 and five individuals in cross 2. Strains: cross 1, NM5970 NM5726 and NM5977; cross 2, NM5953 and NM5853.

### Multiplexed injections

Since every integration plasmid contains both an *FRT* and an *FRT3* site, it is not possible for more than one plasmid to remain integrated in a landing site in the presence of FLP. Thus, co-injection of multiple integration plasmids to increase the efficiency of transgene formation should be feasible. A set of twelve injections sessions were used to demonstrate that the approach is practical. All five coinjections of two plasmids yielded insertions of both, five of six injections of three plasmids yielded insertions of all three, and in a single injection of four plasmids, yielded insertions of three plasmids (Table S2). While this approach can greatly increase the efficiency of transgenesis, unless the individual integration events can be distinguished visually, it is likely more time efficient to inject the targeting plasmids independently than to perform the molecular analysis to distinguish random individual insertions from a multiplexed injection.

### RMCE insertion is highly efficient and accurate

To quantify the efficiency and fidelity of insertions derived using the rRMCE approach, the frequency of insertion of a variety of promoters driving *mNG* or *Scarlet* inserted by rRMCE were tabulated. In 39 injection sessions (Table S2), drug resistant transgenic animals were obtained on 112 of 126 plates (3 P0s/plates) indicating that ~52% of injected animals yield lines (see supplemental methods for calculation). To confirm the fidelity of the method, long range PCR was performed on a set of 26 insertions and the structure of these insertion was confirmed by restriction digestion of the resulting product. In addition, the entire insert sequence of six inserts were confirmed using nanopore technology. These data indicate that the vast majority of insertion events occur by recombination at the expected *FRT* sites. However, a few lines isolated were clearly of unexpected structure. In one case, a plate yielded both animals with TRN Scarlet expression that was specific (*jsSi1864*) and others with slightly brighter TRN Scarlet expression and a broad Scarlet background in all tissues (*jsSi1865*). In another one case, integrations were isolated from a plate that contained animals of two different intensities (*jsSi1955* and *jsSi1955Br*). In both cases, long range PCR across the brighter insertions failed to produce a product, PCR across the dimmer strains showed the expected signal. In another case, a line was isolated which expressed in a pattern consistent with co-integration of both plasmids in a multiplex injection (*jsSi2075*). In one case, only one of two expected FP signals was observed: *jsSi2078* expresses the *myo-2p nls-mNG cis*-marker, but not an expected insert marker. Finally, two insertions isolated were hygR and expressed insert markers but maintained the cyOFP signal of an intact unmodified landing site (*jsSi2081, jsSi2120*). These might represent recombinase-independent integration of plasmids at other sites in the genome. The frequency of these events has not been determined accurately, but they represent a small minority of events recovered. Regardless of their exact frequency, these events are easily distinguished visual from *bona fide* insertions and thus are not a significant impediment to creating rRMCE insertions.

## Discussion

The transgenesis pipeline described here permits the efficient creation of cis-marked single copy insertions in eight days. A multitude of vectors and landing sites permits the creation of insertions marked with distinct FPs, expressed in distinct tissues, and on different chromosomes thus providing great flexibility in the design of more complex transgenic strains that require multiple insertions. In addition, the method is efficient enough that multiplex injections can be used to triple or quadruple the efficiency of transgene generation in cases where the individual insertions can readily be distinguished. The interactions observed between transcription units have been well characterized to provide guidance to the user in the design of transgenic lines. To mitigate the potential reduction in utility of transgenes with unexpected cross transcription unit interactions, germline Cre expressing lines are also described that permit the modification of insertions by highly efficient Cre mediated excision using simple crosses. Finally, a variety of reagents are described that facilitate the customization of both landing sites and integration vectors to extend the approach.

### Design considerations

How to organize the structure of the landing sites and integration vectors was considered at length. One approach is to use a split-selectable marker approach (Stevenson et al., 2020; El Mouridi et al., 2022) which has been favored because it essentially eliminates the background of extrachromosomal arrays by requiring recombination between the landing site and targeting vector to obtain resistance. However, since the RMCE system utilized herein expresses FLP constitutively, background is essentially eliminated even without splitting the selection marker. The major disadvantage of a split selectable marker is that each landing site is restricted to a specific selection marker. By contrast, rRMCE system landing sites can be used with any selectable marker including *unc-119*. Thus, a split drug resistance marker was not adopted, though derivatives that use this approach could easily be created with one cloning step and one RMCE step.

The landing sites developed herein harbor a cell specific promoter driving cyOFP in the landing site and the approach uses the recombination event to swap the FP associated with the promoter which provides an easily method for confirming the position of the integration event. The major advantage of the approach used herein is that the cloning protocol is much simpler without having to insert both a promoter and FP for detection into the integration vector in addition to expression cassette of interest. The approach chosen favors simplifying the cloning process at the cost of the initial creation of more landing sites. It is debatable as to which is the most time and resource efficient approach. Simplifying the cloning process was chosen since most labs focus on a select set of cell types. However, a lab that works in many different tissues might opt for a distinct approach. In line with this, landing sites in which the recombination site is placed upstream of the promoter were also constructed. Although these were intended for constructing insertions that do not express a visible cis-linked marker, they could be used for to integrate plasmids that contain a cell-specific promoter and FP, a selectable marker, and an expression cassette of choice without the construction of new landing sites.

The *loxP* site in the integration plasmids was positioned such that both the drug/marker selection transcription unit and the *cis*-promoter FP marker could be simultaneously excised for the insertion using Cre recombinase. However, as currently implemented, it is not possible to excise the selection cassette without excising the *cis*-promoter FP linked to the insertion. An additional disadvantage of the approach chosen is that insertions that retain the *cis* marker and selection cassettes are potentially incompatible with experiments using *loxP*/Cre as a tool to conditionally alter gene function. To mitigate this concern, the pHyg8 series vectors that lack a *loxP* site were created. Insertions made using these vectors lack a second *loxP* site and are completely resistant to Cre recombinase as evidenced by the germline Cre expressing strains created to simplify excision of the floxed markers of standard rRMCE insertions. The pHyg8 series vectors could also be modifying by adding *lox2722* or *lox511* sites flanking the HygR gene which would permit excision the selection cassette independently of the *cis* FP marker.

### Crosstalk in insertions containing multiple transcription units

The transgenic animals created using rRMCE typically contain at least three distinct transcription units: a tissue specific promoter driving a FP, a ubiquitous promoter expressing a drug resistance gene, and a transcription unit unique to the specific insertion. In multiple insertions characterized crosstalk between these transcription units was detected. The *C. elegans* genome lacks the classic insulator protein CTCF (Heger et al., 2009) and studies have argued that the main influence on gene expression specificity in *C. elegans* is the distance between enhancers and promoter (Quintero-Cadena and Sternberg, 2016). Operons are also common in *C. elegans* (Blumenthal and Gleason, 2003) and recent work suggests that polycistronic transcription is more prevalent than previously thought (Bellush and Whitehouse, 2019). The crosstalk observed in rRMCE transgenes could in most cases logically be explained either by polycistronic transcription resulting from failure of RNA polymerase II termination or by enhancer sharing. However, in the case of crosstalk between the *myo-2p* and *myo-3* promoter (Figure 2C), the downstream promoter is co-opting the upstream promoter which is inconsistent with a polycistronic model. Thus, it is reasonable to presume that both enhancer sharing and polycistronic transcription contribute to the observed crosstalk.

Unfortunately, since some promoters exhibit sensitivity to crosstalk and others are resistant, no simple transgene design paradigm is likely to eliminate crosstalk in all cases. In C. elegans, transcriptional termination is inefficient often occurring over a region of over a kb after the polyadenylation site (Haenni et al., 2009) and failure of termination has been shown to greatly increase expression of adjacent transcription units (Miki et al., 2017). Thus, increasing spacing between transcription units and using specific relative orientations of transcription units is likely to minimize or eliminate crosstalk in most cases. The orientation (*myo-2p* FP > <rps0-hygR goi-p FP>) was largely devoid of crosstalk and likely represents the most logical order to use for most transgenes. Flipping the orientation of the *goi-p* FP may further reduce crosstalk, but at the expense of potential influence on the goi-p from sequences in the genome adjacent to the insert as was observed for the Chr IV insertions (Figure 5F, H, L, Figure S4F, H, Figure S6J). In addition, matching promoters with appropriate 3’ UTRs may also help elevate crosstalk due to termination failure. Termination in *C. elegans* occurs by two distinct mechanisms (Miki et al., 2017). Importantly, whether termination is *xrn-2* dependent or independent is dictated by the promoter and also requires a compatible polyadenylation downstream region (Miki et al., 2017). The commonly used *act-4* and *tbb-2* 3’ UTR are *xrn-2* dependent and thus may not be appropriate for use with genes using an *xrn-2* independent termination mechanism. Rather the *xrn-2* independent *unc-54* 3’ UTR may be more appropriate. The most appropriate approach might be to expend the effort to use the native 3’ UTR when creating cell specific promoter constructs whose utility requires an expression pattern that matches the in vivo setting. It should also be noted that it is unclear whether the existing resistance cassette in rRMCE vectors uses matching promoters and 3’ UTRS. The hygR cassette uses the unc-54 3’ UTR. However, most, but not all, ribosomal protein genes are xrn-2 dependent (though the state of *rps-0* was not defined in Miki et al., 2017). The fact that transgenes that have hygR oriented in the same orientation as the goi-1p often show severe cross talk be the result of promoter-3’UTR mismatch and potential ameliorated by changing either the promoter or 3’ UTR.

### Error rates

To assess the fidelity of rRMCE insertion, over 30 insertions were analyzed by either PCR or nanopore sequence analysis (Table S4). All the PCR amplified insertions were of the expected size and restriction analysis of the products yielded the expected digestion pattern and all the sequenced insertion were identical to the expected sequence indicating that the technique is very reliable. However, several insertions that were not of the expected structure were isolated during this study. All of these were visually identified during the isolation of hygR resistant insert clones. Typically, insertion bearing homozygous animals were isolated on day 8 after injection. Often, these plates contained hundreds of healthy L3 to adult staged animals, approximately one third of which were identifiable as homozygous animals based on FP marker expression. The vast majority of these animals showed the expected insert fluorescent expression pattern. However, occasionally, animals with unusual patterns were also observed and isolated. The interpretation of data regarding these insertions is based upon RMCE insertions occurring as proposed in Nonet (2020). Specifically, that extrachromosomal arrays of the input plasmids are formed in the cytosolic P0 germline and that subsequently these arrays enter the nucleus after fertilization and are then broken down by FLP-mediated recombination in the F1 germline. These monomers or small concatemer array fragments are subsequently integrated into landing sites and then resolved (either fixed in the genome or re-excised) by additional FLP recombination leaving a single copy insertion (see Figure 7 A-C of Nonet, 2020).

**Figure 7.**
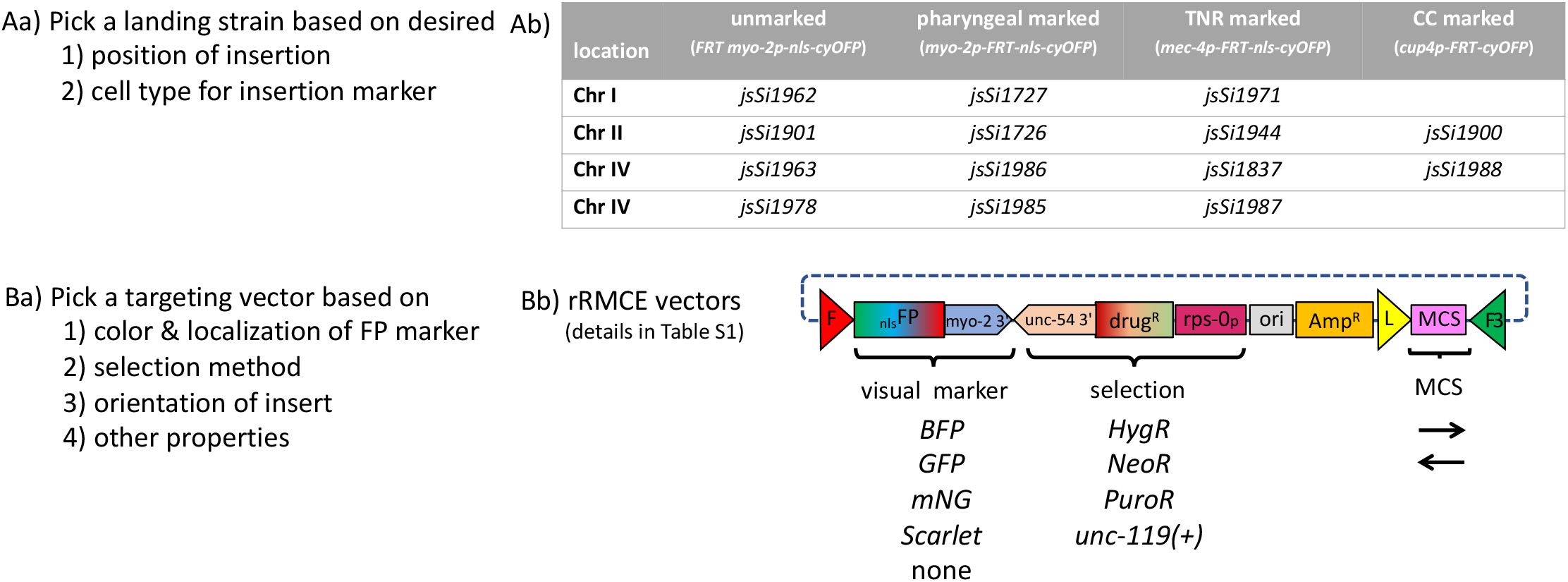
Design options for rRMCE transgene construction. Transgenes created using rRMCE are integrated at specific genomic positions and express a fluorescent protein marker in a specific cell-type dependent on which combination of integration vector and landing site are utilized. **Aa)** Factors to consider in deciding on a landing site. One should take into consideration both where in the genome one would like the transgene to reside and how one would like the transgene to be marked. **Ab)** A table of the available landing sites. These are available on four chromosomes and are marked with either a pharyngeal *myo-2p*, a TRN-expressing *mec-4p*, or a coelomocyte-expressing *cup-4p*. In addition, a set of landing sites with the FRT site upstream of the *myo-2p* allows for the creation of unmarked lines. The unmarked lines are also available in an *unc-119(ed3*) background (see Table S4). In considering which landing site to use one should be aware that the TRN and coelomocyte promoters are weaker than the *myo-2p* and the marker signal observed in lines integrated at these landing sites is more difficult to detect in the dissecting microscope, though all are easily detected at 20X using a compound microscope. **Ba)** Factors one should consider in deciding on an integration vector to use to clone your sequences of interest. These include the color and the subcellular localization of the fluorescent visual marker, the type of selection method, and the orientation of both the selection marker and the insertion. Vectors with cytosolic fluorescent markers yield stronger signals than vectors with nuclear localized fluorescent markers. Vectors lacking a *loxP* site which render the insertion non-excisable by Cre and vectors which exclude the backbone sequences from integrating are also available. **Bb)** Schematic diagram of the organization of rRMCE vectors showing the options for the visual marker, the selection method and the orientation of the insert. Most visual marker options are available with HygR as the selection method, but the variety for others selection methods is very limited. See Table S1 for a detailed list of available vectors. The vectors (except the cytosolic Scarlet RR series) do not contain BsaI sites and thus can relatively easily be modified using Golden Gate cloning methods analogous to those used to create the HygR series of vectors.

The first class of animals observed were animals with a brighter expression pattern (i.e., *jsSi1955Br*). When these animals were backcrossed to animals carrying an unmodified FLP expressing landing site, the trans-heterozygotes segregated both bright and dim animals. PCR across the insertion failed in the bright animals and was successful in the dim animals suggesting that these insertions are trapped intermediates that contain more than one concatenated insertion plasmid. Since the source of FLP is removed during the cassette exchange, the timing of the exchange may result in insertions that still contain multiple copies of the insert but now lack the FLP enzyme required to resolve the insertion by recombination between FRT or FRT3 sites of adjacent insert copies of the targeting plasmid. Isolating heterozygote rather than homozygous animals on day 8 may minimize the trapping of intermediates by providing an additional generation in the presence of FLP for excision to take place.

A second class of insertions were occasionally observed in multiplex injections (i.e., *jsSi1075*). These displayed the expression pattern of multiple distinct input plasmids and often also showed broader non-specific expression of FP in many somatic tissues. Outcrossing of these animals to animals carrying the unmodified FLP expressing landing site did not alter the transgene expression profile. However, outcrossing through a Cre expressing strain eliminated the broad FP expression and reduced the FP expression to that consistent with a single insertion plasmid. Based on these data, these insertions likely represent insertion of fragments of multiple plasmids that recombined non-homologously during array formation in the P0 animal. The broad expression pattern likely represents crosstalk between hygR cassettes of the insertion and adjacent insert sequences.

Another class of insertion observed were homozygous insertions that still contained the marker fluorescent pattern of the unmodified landing site (i.e., *jsSi2021*). These types of insertions were isolated from plates that on day 8 only contained ‘heterozygous” hygR insertions that still expressed cyOFP from an unmodified landing site. However, some of these animals segregated 100% animals expressing the *goi-p* FP expression pattern indicating they were homozygous for an insertion, but they failed to segregate animals that switched the cyOFP *cis*-marker. PCR confirmed the landing sites were unmodified. These insertions are likely either FLP-mediated insertions of the plasmid at pseudo-*FRT* sites or random recombinase-independent insertions. However, no other molecular or genetic analysis was performed to determine the structure or position of these insertions.

It is difficult to assess the exact frequency of the events described above during rRMCE. First, it is difficult to assess the number of independent insertions occurring on an injection plate, especially when hundreds of different hygR F2 animals are present. Second, most injection plates with resistant animals were not screened exhaustively for unusual patterns. On day 8, animals with very strong mosaic FP expression patterns were often still present on the drug selection plates along with other unhealthy animals. Only healthy L4 animals were picked to establish insertion lines. In cases, where live animals were present on the plate, but no healthy homozygotes could be identified, either healthy heterozygotes were isolated, or the plate was re-screened several days later. In cases where the targeting plasmid expresses a visual marker, unusual events such as those described above can largely be avoided by choosing the animals that express the common expression pattern observed on several different injection plates. However, in cases where the insert does not yield a visible signal, molecular analysis of the insert can readily confirm the structure of the insertion.

### Comparison of methods

There are now multiple different approaches available to create single copy transgenic animals (Praitis et al., 2001; Frokjaer-Jensen et al., 2008; Kim et al., 2014; Dickinson et al., 2015; Paix et al., 2015; Silva-García et al., 2019; Ghanta and Mello, 2020; Nonet, 2020; Stevenson et al., 2020; El Mouridi et al., 2022; Yang et al., 2022). rRMCE provides a time efficient method to create single copy transgenes that are cis-marked to simplify tracking the insertion in crosses. Creating such transgenes using other common insertion methods is more complicated because the cis-marker transcription unit must be co-inserted with the goi transcription unit while it is an integral component of rRMCE. RMCE in general, and rRMCE specifically is also relatively insert size independent. The major limiting factor is how large an insert can be integrated is the size of the targeting plasmid that can be constructed; roughly 20-25 kb for the pHyg vectors described herein. rRMCE also requires minimal hands-on time and has high enough fidelity that initiating experimental approaches using newly developed transgenic lines without molecular characterization is not a significant risk. rRMCE should be particularly useful when multiple distinct transgenes need to be combined to create a strain of interest.

### Tools for further development of RMCE methods

rRMCE provides a time and resource efficient way to create transgenic animals. However, the design features emphasized in this implementation of the approach may not be convenient, compatible, or favored by other scientists. For those who find the efficiency and fidelity of the approach appealing, but the specific implementation problematic, several tools were derived herein to facilitate modification and further refinement of the method. First, several tools were created to simplify the creation of novel landing sites. Intermediate landing sites were created that permit one to design and introduce landing sites with novel organization with a single cloning step followed by a single RMCE integration. Second, a SEC-landing site cassette which requires only the addition of two targeting homology arms should facilitate introduction of landing sites at completely novel genomic location using an SEC-CRISPR approach. Second, most of the vectors described for rRMCE were constructed using Golden Gate cloning and are devoid of BsaI sites allowing them to be easily modified via Golden Gate cloning methods. It is hoped that these tools will stimulate the development of other creative uses for RMCE in the worm.

## Abbreviations

MosSCI: Mos Single Copy Insertion
TRN: touch receptor neuron
CRISPR: Clustered Regularly Interspace Short Palindromic Sequences
RMCE: Recombination-Mediated Cassette Exchange
rRMCE: rapid RMCE
3’ UTR: three prime untranslated region
mNG: monomeric Neon Green FP
FP: Fluorescent Protein
GFP: Green FP
BFP: Blue FP
cyOFP: cyan-excitable Orange FP
SEC: Self-Excision Cassette

## Acknowledgements

I thank Emma Knoebel for performing some of the injections in the last few months of this project, Matt Rich, Adam Hefel, and Erik Jorgensen for providing a *C. elegans* codon optimized *Cre* gene, Jordan Ward for comments on the manuscript, and Tim Schedl for discussions during development of this methodology. I apologize to those who have published methods for C. elegans genome manipulation that I did not cite due to space restrictions. This work was funded in part by R01GM14168802.

## Methods

### Nomenclature

*C. elegans* RMCE insertions into a landing site locus (e.g., *jsSi1726*) should technically be called *jsSi1726 jsSi#* according to *C. elegans* nomenclature rules (Tuli et al., 2018), but were referred to in the paper as *jsSi#*.

This work clearly demonstrates that the orientation of inserts is a major determinant of expression profiles in transgenes. However, the current *C. elegans* nomenclature system for transgenes does not address orientation of insertions. To deal with this issue this paper incorporates a nomenclature system in which insertions that are in opposite orientation of that implied by the general nomenclature system are surrounded by <{region reverse oriented} to indicate that this section of the insertion is in the opposite orientation. This nomenclature system is used in Tables S3 and S4 which describes the transgenes created in this work.

### *C. elegans* strain maintenance

*C. elegans* was grown on NGM plates seeded with *E. coli* OP50 on 6 cm plates. Stocks strains were maintained at RT (~ 22.5°C). RMCE experiments were performed at 25°C.

### Plasmid microinjections

Injections were performed as described in Nonet (2020) except that a paint brush (Robert Simmons E51 liner 10/0) was typically used to mount animals onto the agar pad before injections. This modification increased the injection survival rate to >90% of animals. Most animals were only injected in a single gonad. DNAs were injected at ~ 50 μg/ml in 10 mM Tris pH 8.0, 0.1 mM EDTA. Specific DNA concentrations for the 39 injection sessions used to assess integration efficiency are found in Table S2. Isolation of RMCE insertions Landing sites we constructed using the RMCE *sqt-1* Rol screening strategy and performed as outlined in Nonet, 2020. For rapid RMCE using drug selection, injected P0 animals were pooled 3 per plate. For drug selection, the following were added directly to worm plates 3 days after injection: *HygR* selection −100 ul of 20 mg/ml Hygromycin (HygroGold™ InvivoGen, San Diego, CA) or Hygromycin B, GoldBio, St. Louis, MO), *NeoR* selection - 500 ul of 25 mg/ml G418 disulfate (Sigma, St. Louis, MO), *PuroR* selection - 500 ul of a solution of 10 mg/ml Puromycin dihydrochloride (GoldBio), 0.1% Triton X100 (Sigma). Eight days after injection L4 animals homozygous for insertions could usually be identified on injection plates. In some cases, only a few heterozygous *hygR* animals were present, in which case, homozygous animals were isolated a few days later. The integration plasmid and parental strain used to construct each novel transgenic insertion are listed in Table S3 and the strains containing these alleles and all other strains used in the study are listed in Table S4.

### Characterization of RMCE landing sites and insertions

The structure of all novel RMCE landing sites and some RMCE insertions were analyzed using long range PCR of genomic DNA as previously described (Nonet, 2020). Oligonucleotide NMo6563/6564 were used to amplify Chr I insertions, NMo3880/3884 or NMo3887/3888 for Chr II insertions, NMo3889/3890 for chr IV insertions, and NMo7280/7281 for Chr V insertions. The sequence of all landing sites was confirmed by nanopore sequencing of PCR products spanning the insertion. Some RMCE insertions into landing sites were sequenced, but most were only analyzed by restriction digestion of PCR products to confirm the insertion structure. See Table S4 for specifics.

### Plasmid and vector constructions

The term vector, rather than plasmid, was used to distinguish the parental integration vectors from integration plasmids that contain specific sequences inserted in the MCS of the vectors. All PCR amplifications for plasmid and vector constructions were performed using Q5 polymerase (New England Biolabs, Ipswich, MA). Most PCR reactions were performed using the following conditions: 98°C for 0:30, followed by 30 cycles of 98°C for 0:10, 62°C for 0:30, 72°C for 1:00/kb). PCR products were digested with DpnI to remove template if amplified from a plasmid, then purified using a standard Monarch (New England Biolabs) column purification procedure. Restriction enzymes (except for LguI), T4 DNA ligase, and polynucleotide kinase were purchased from New England Biolabs. Golden Gate (GG) reactions (Engler et al., 2008) were performed as described in Nonet, 2020 except that in some cases LguI (Thermo Scientific™, Waltam, MA) was used in place of SapI. The *E. coli* strain DH5α was used for all transformations. Sanger sequencing was performed by GENEWIZ (South Plainfield, NJ) and nanopore sequencing by Plasmidsaurus (Eugene, OR). Oligonucleotides were obtained from MilliporeSigma (Burlington, MA) and synthetic DNA fragment were purchased from Twist Biosciences (South San Francisco, CA). A detailed description of all constructs is provided in supplemental methods. The sequence of all plasmids, vectors and synthetic fragments is provided in Table S5 and oligonucleotides used in the study are provided in Table S6.

### Microscopy

Screening of worms for fluorescence during the RMCE protocol was performed on a Leica (Heerbrugg, Switzerland) MZ16F FluoCombi III microscope with a planapo 1X and planapo 5X LWD objective for high power observation illuminated using a Lumencor (Beaverton, OR) Sola light source.

For imaging or quantification of fluorescence, worms were mounted on 2 % agarose pads in a 2 μl drop of 1 mM levamisole in phosphate buffered saline. 10 to 20 L4 animals were typically placed on a single slide. Animals were imaged using a 10X air (na 0.45) or 40x air (na 0.75) lens on an Olympus (Center Valley, PA) BX-60 microscope equipped with a Qimaging (Surrey, BC Canada) Retiga EXi monochrome CCD camera, a Lumencor AURA LED light source, Semrock (Rochester, NY) GFP-3035B and mCherry-A-000 filter sets, a Chroma (Bellows Falls, VT) 89402 multi-pass filter set and a Tofra (Palo Alto, CA) focus drive, run using Micro-Manager 2.0ß software (Edelstein et al., 2014). For image acquisition, the 485 and 560 nm LEDs were at full power, 200 ms or 500 ms exposure were used for 10X images and a 500 ms exposure was used for 40X images. The size of L4s often necessitated taking images with the animal in a diagonal position across the field. For figure panels such images were rotated leading to the presence of black corners in many panels. These black corners were adjusted to the mean intensity of a 3-pixel width line adjacent to the diagonal of each corner using a custom imageJ macro. Images of cyOFP expressing animals were taken with the 485 nm LED and a multi-pass filter set and pseudo-colored orange. Long exposure images were created by reducing the window level 7.5-fold compared to short exposure image.

## Data Availability

Critical worm strain and plasmids will be made available at the Caenorhabditis Genetics Center and at Addgene. All other reagents are available upon request to Michael Nonet.

## Supplemental Materials

Sup. Tables, Figures, Legends and Methods available at https://sites.wustl.edu/nonetlab/manuscripts/

Table S1. Table of rRMCE vectors.

Table S2. List of injections used for quantifying integration frequency.

Table S3. List of alleles and transgenes used in this study.

Table S4. List of C. elegans strains used in this study.

Table S5. List of DNA constructs used in thus study. Table S6. List of oligonucleotides used in this study. Figure S1. Overview of original RMCE method. Figure S2. Strategy for creating rRMCE landing sites. Figure S3. Construction of rRMCE vectors.

Figure S4. Landing site effects on specificity of expression in rRMCE transgenes

Figure S5. Overview of the rRMCE method for TRN, coelomocyte, unmarked, and *unc-119*(+) lines Figure S6. Additional TRN, coelomocyte, and unmarked lines

Figure S7. Methodology for constructing novel rapid RMCE landing sites

Supplemental methods.

Supplemental figure legend.

