## Supplemental Figures and Legends for "Rapid generation of *C. elegans* single copy transgenes combining RMCE and drug selection"

### Overview of original RMCE method

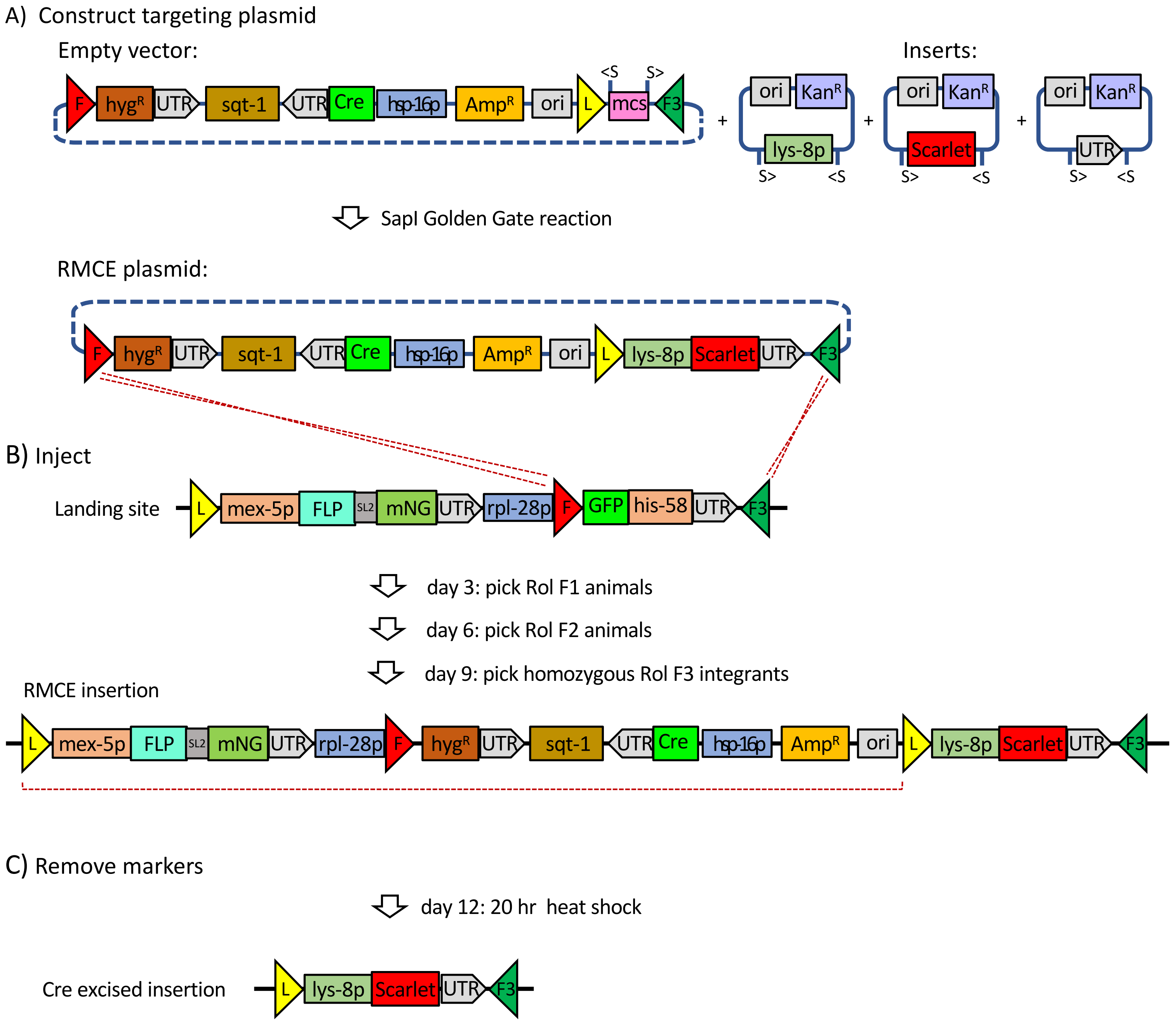


Figure S1

Construction of new landing sites

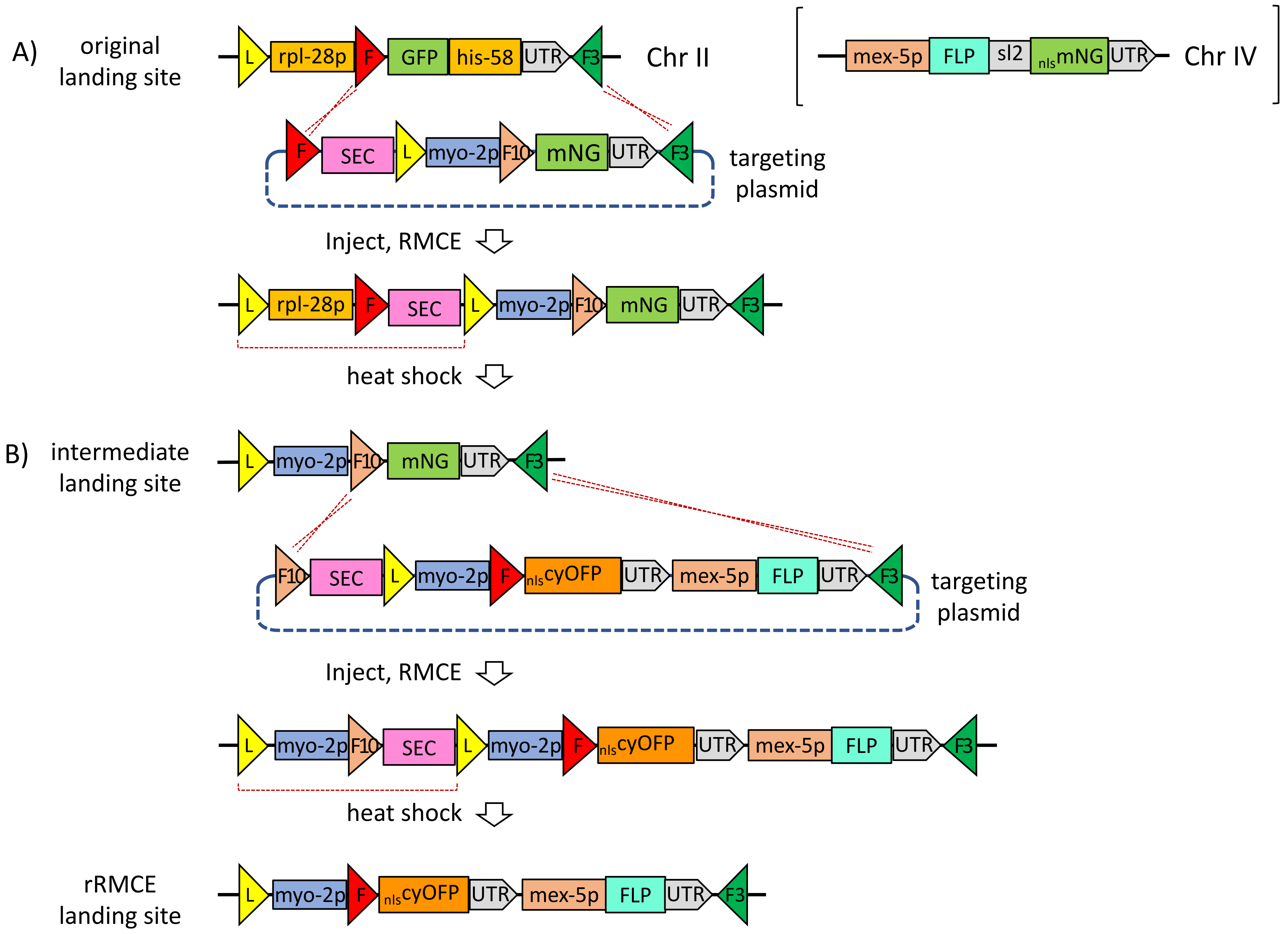


Figure S2

Construction of new RMCE vectors

A) Assembly of pHygG1

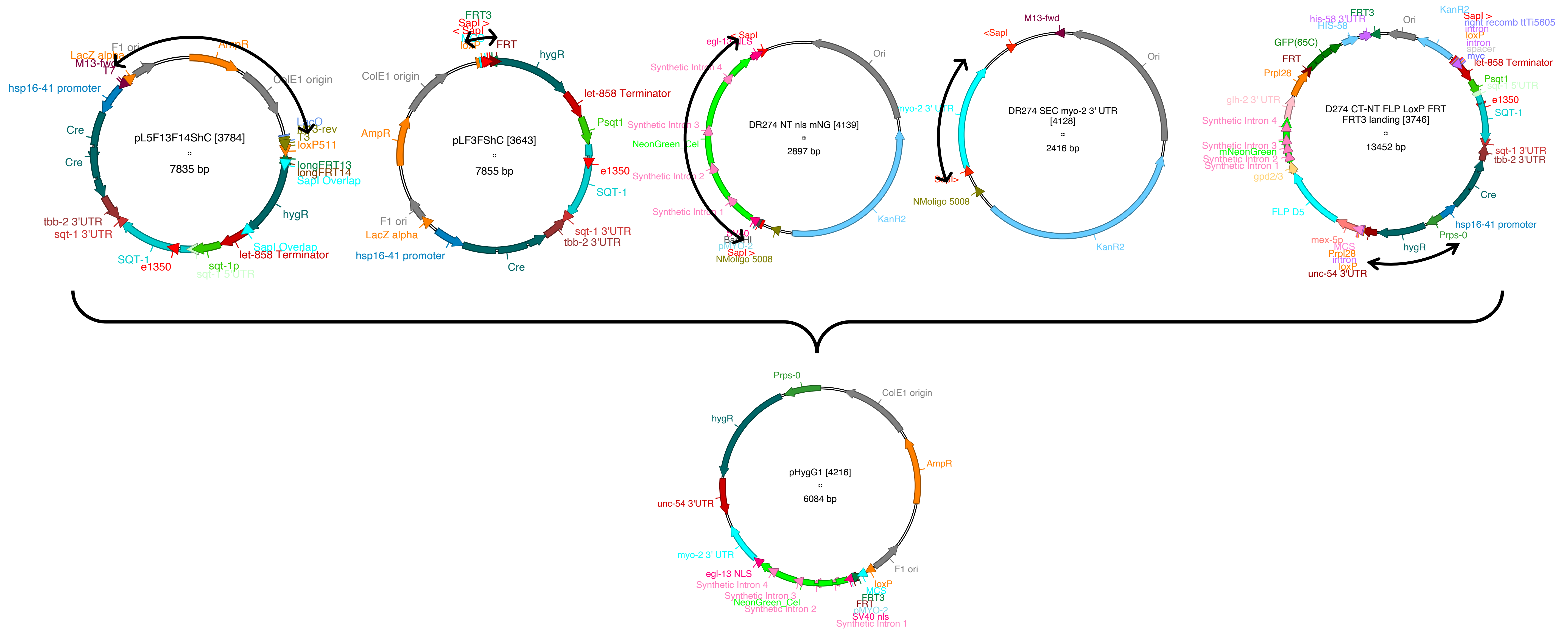


B) Flow Diagram of rRMCE vector construction

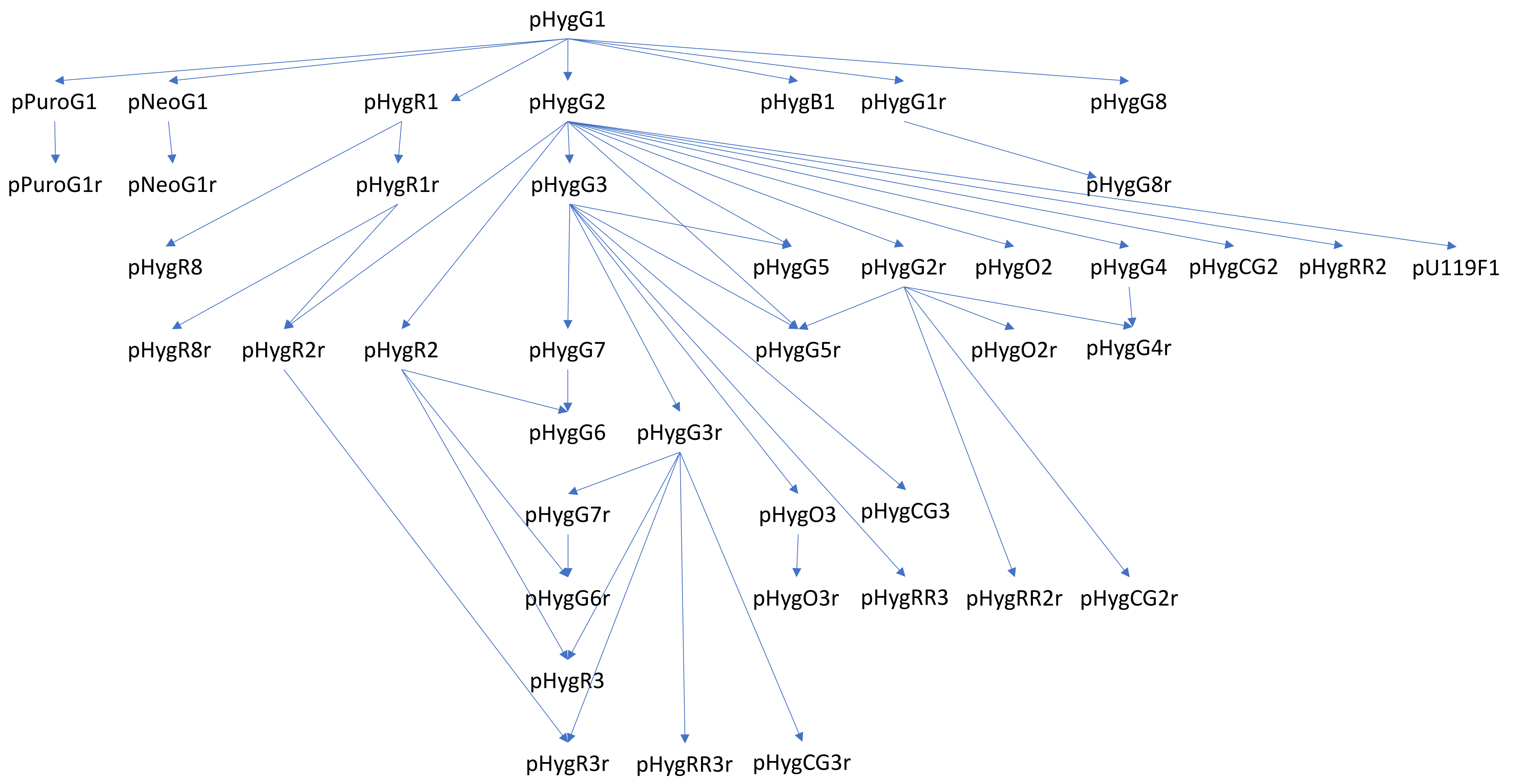


Figure S3

Orientation effects on specificity of expression at distinct landing sites

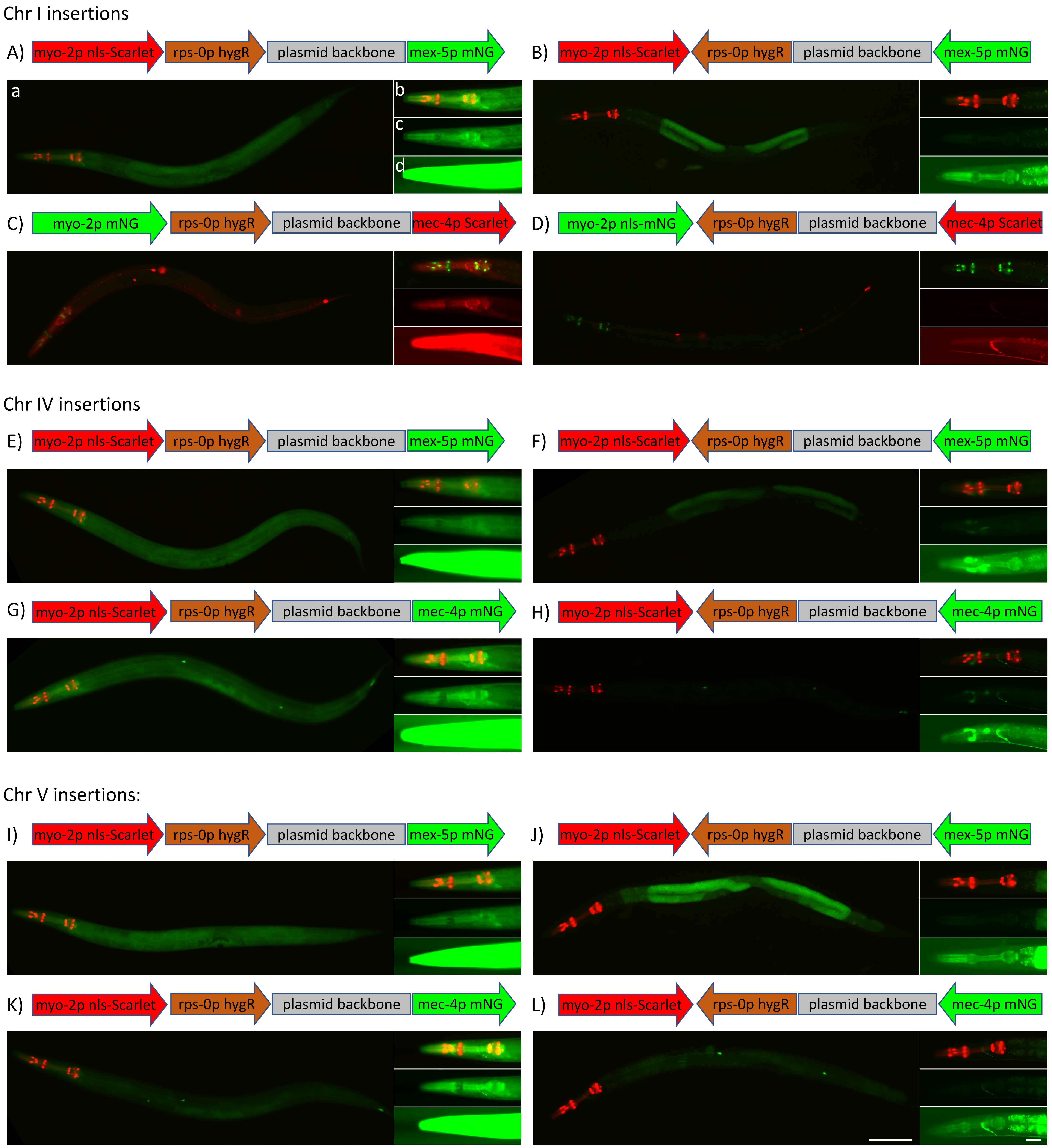


Figure S4

Overview of rRMCE method applied to create TRN, coelomocyte, unmarked, and unc-119 marked lines

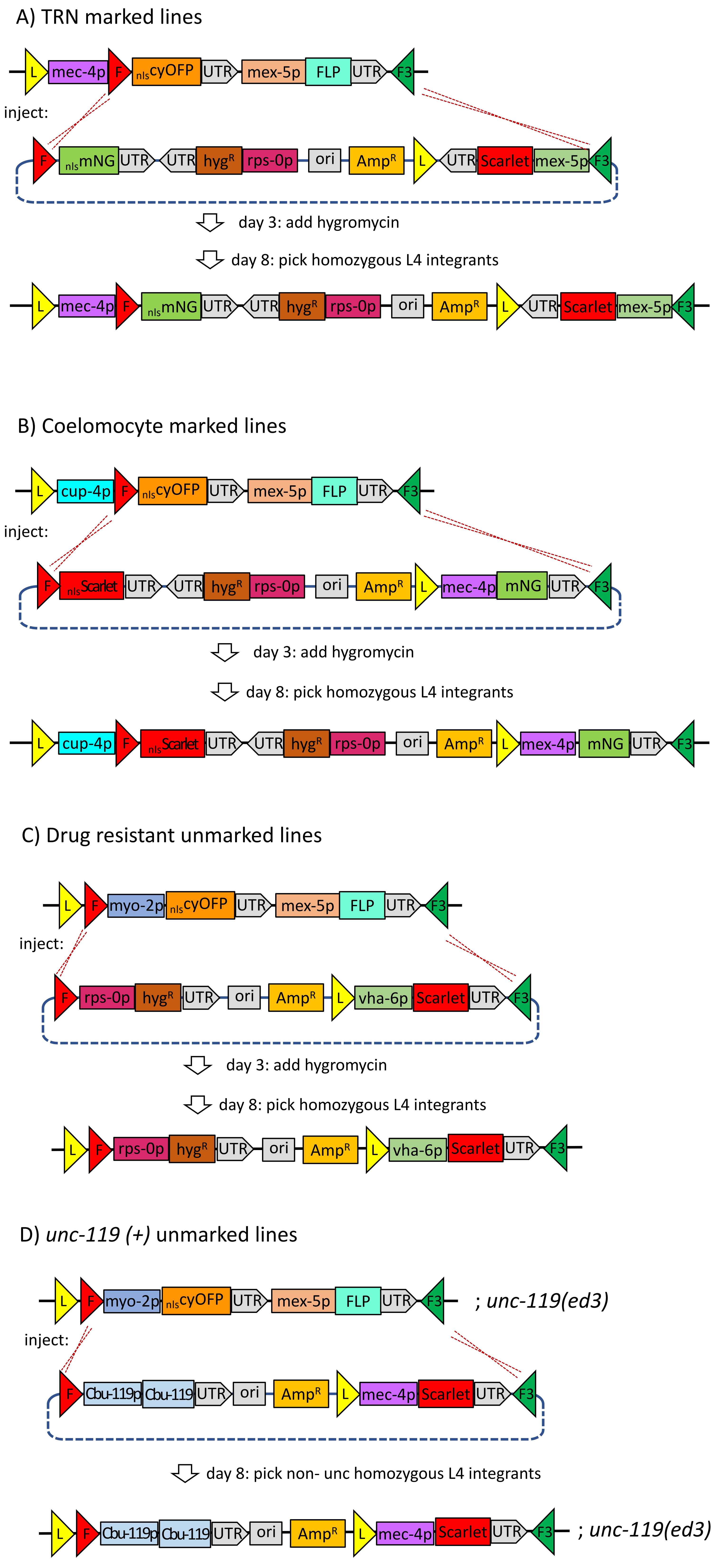


Figure S5

Additional exampled of phagyngeal, TRN, coelomocyte, and unmarked lines

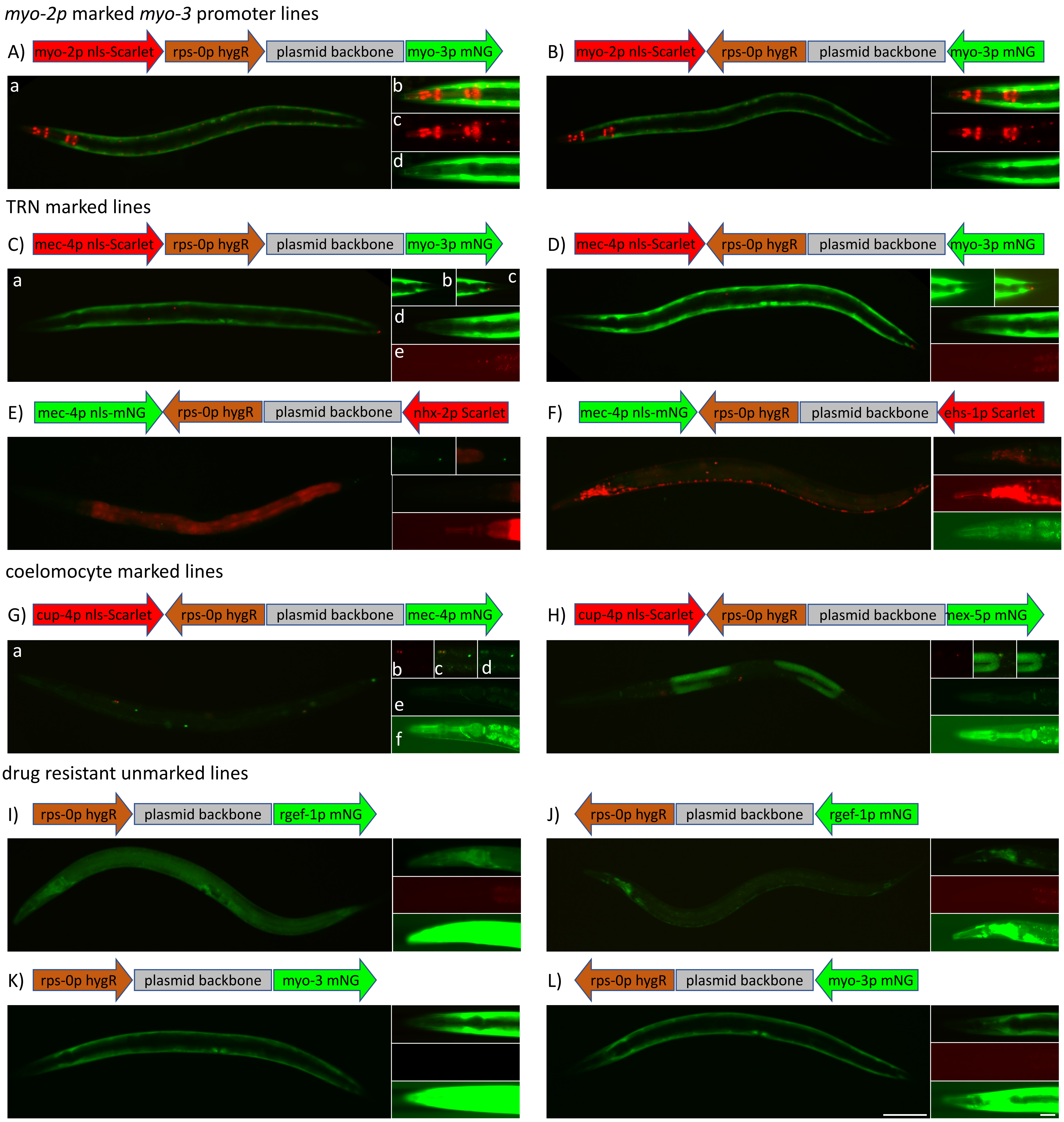


Figure S6

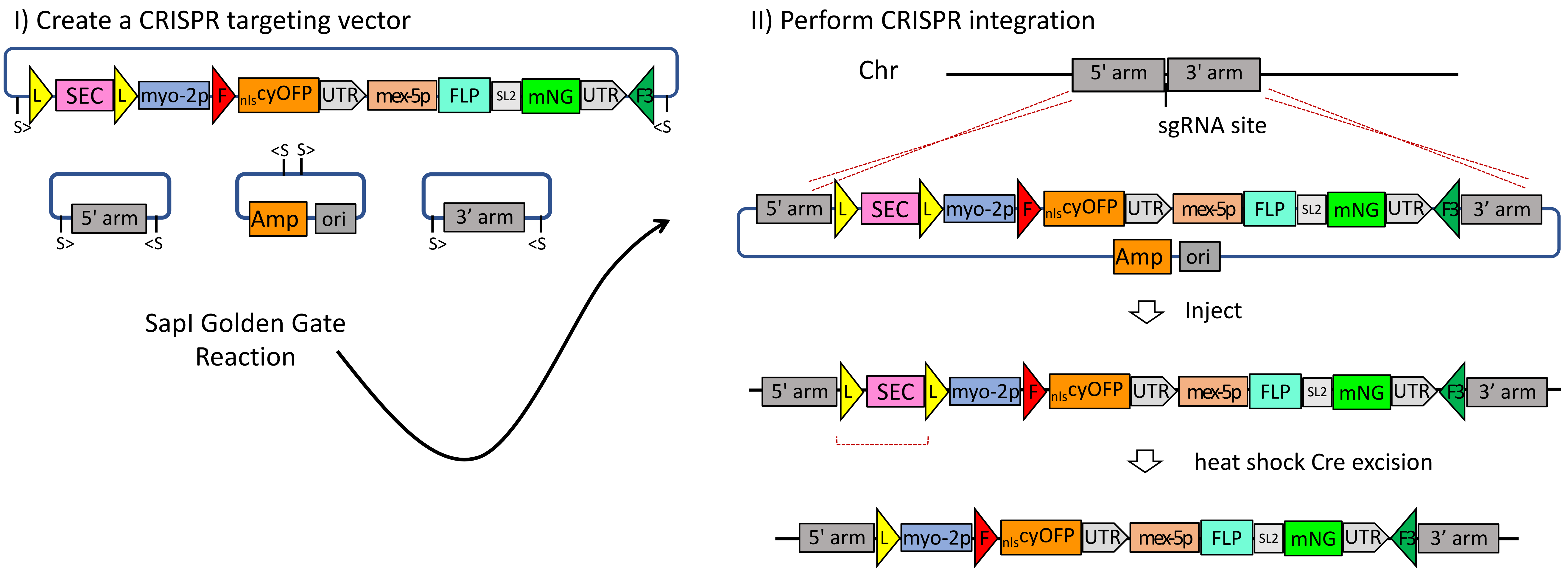

I) Create a CRISPR targeting vector
II) Perform CRISPR integration
Chr
5' arm
3' arm
sgRNA site
L
SEC
myo-2p
F
nlscyOFP
UTR
mex-5p
FLP
SL2
mNG
F3
S>
<S
Amp
ori
SapI Golden Gate Reaction
Inject
heat shock Cre excision


Figure S7

**Supplementary Figure Legends**

Figure S1. Overview of the original RMCE method

**A)** A targeting plasmid is constructed using a *SapI* Golden Gate assembly reaction using the pLF3FShC targeting vector and clones encoding the *lys-8p*, the fluorescent protein *Scarlet* and the *tbb-2* 3' UTR. S symbols represent *SapI* sites and > symbols indicate the orientation of sites in the plasmids. Colored triangles represent *loxP* (yellow), *FRT* (red) and *FRT3* (green) recombinase sites.

**B)** The resulting targeting plasmid is injected into adult animals of a landing site strain. This example shows a 'single component' landing site with a cassette containing the *mex-5p* driving Cre in the landing site. However, other landing sites, called 'two component' landing sites, do not contain the cassette and must be paired with the bqSi711 IV *mex-5p* FLP transgene {Macías-León and Askjaer, 2018, #78303; Nonet, 2020, #20343}. The landing site strain expresses nuclear mNG in all cells and FLP in the germline. The injected animals (3 animals/plate) are incubated at 25°C. On day 3, Rol animals are picked (5 animas/plate). On day 6 the F1 plates are screened for rare F2 Rol animals which are almost always integration events. On day 9, homozygous GFP(-) animals are cloned.

**C)** On day 12 L1 homozygous animals are heat shocked to excise the SEC (Self Excisable Cassette) to identify non-Rol animals harboring the final insertion.

Figure S2. Strategy for creating rRMCE landing sites

**A)** The structure of a Chr II RMCE landing site (*jsSi1579*) is depicted along with the structure of the FLP expressing insertion *bqSi711*. A strain containing both insertions was injected with a *myo-2p FRT10 mNG* targeting plasmid which yielded via RMCE integration a homozygous *sqt-1*(Rol) *hygR myo-2p mNG* strain. That strain was subsequently heat-shocked to remove the *sqt-1*(Rol) *hygR* Self-Excising Cassette which yielded the homozygous non-Rol *myo-2p* mNG intermediate landing site strain.

**B)** The intermediate *loxP FRT10 FRT3* landing strain (also harboring the *bqSi711* FLP D5 transgene) was injected with a *myo-2p FRT* nls-*cyOFP mex-5p FLP* targeting plasmid which yielded via RMCE integration a homozygous *sqt-1*(Rol) *hygR myo-2p nls-cyOFP* strain. This strain was subsequently heat-shocked to remove the *sqt-1*(Rol) *hygR* Self-Excising Cassette which produced the desired *myo-2p nls-cyOFP mex-5p* FLP landing site strain. The structure of this and other landing sites created herein was confirmed by PCR amplification across the insertion site followed by nanopore sequencing of the product.

Figure S3. Construction of the rRMCE vectors.

**A)** Diagram of the assembly of pHyg1 from 5 fragments derived from existing plasmids. The pHyg vector constitute a major reorganization of the vector design compared to the original pLF3FShC vector which formed the basis of the original RMCE method. The key differences are 1) hygR now is driven by the

*rps-0p* while in the original vector it was promoter-less (and then driven by the *rps-28* promoter in the landing site after insertion thus providing selection for insertion). 2) In the original vectors the plasmid backbone was not incorporated into the final insertion, while in the new method it is. To accomplish this, the position of the *loxP* site was moved relative to the backbone. 3) No fluorescent reporter was present in the original vector. Insertions were screened for by loss of fluorescence derived from the landing site. In contrast, most of the new pHyg vectors harbor a fluorescent marker. 4) The original vectors contained multiple BsaI sites. The vast majority of new vectors were assembled using a BsaI Golden Gate strategy and thus eliminate all BsaI sites (except the cytosolic Scarlet RR series). The new pHyg vectors were assembled from plasmids that had previously been modified to remove BsaI sites.
**B)** Flow diagram showing the relationship between various pHyg vectors and derivatives described in Table S1. Note that not all vectors were used in this study, and some have never been formally tested for function. However, the modifications that have been made are minor and are highly unlikely to affect functionality for RMCE. Precise descriptions of the modifications made for all steps shown in this flow diagram are found in Supplemental Methods.

<u>Figure S4. Landing site effects on specificity of expression in rRMCE transgenes</u>
**A-L)** Effects of orientation on expression of *mex-5p* mNG reporter and *mec-4p* Scarlet reporter *myo-2p* marked insertions on different chromosomes.
Images of representative L4-staged rRMCE insertion transgenic animals. Shown on top of each set of images is a diagram of the insertion depicting each transcription unit and the plasmid backbone (to scale). **a)** Widefield view of an L4 animal showing mNG (green) and Scarlet (red) fluorescent signals. **b-d)** High magnification view of the pharynx of another representative individual: (b) a merge of mNG and Scarlet signals, (c) the reporter signal (d) and a long exposure of the reporter signal.
**A-D)** Chr I insertions. **E-H)** Chr IV insertions. **I-L)** Chr V insertions.
Note the unexpected GFP(+) cells adjacent to the anterior pharynx in panels F and H. Strains shown: A) NM6093 *jsSi2128*, B) NM6053 *jsSi2048*, C) NM6073 *jsSi2079*, D) NM6111 *jsSi2073*, E) NM6098 *jsSi2097*, F) NM6089 *jsSi2088*, G) NM6097 *jsSi2096*, H) NM6108 *jsSi2091*, I) NM6091 *jsS2106*, J) NM6112 *jsSi2101*, K) NM6101 *jsSi2108*, L) NM6148 *jsSi2153.* Scale Bars: 100 µm (whole animal images) and 25 µm (pharyngeal image).

<u>Figure S5. Overview of the rRMCE method as applied to TRN, coelomocytes, unmarked, and *unc-119*(+) lines</u>
The structure of a landing site, an integration plasmid and the final insertion are shown for creation of **A)** the TRN marked line shown in figure 5B, **B)** the coelomocyte marked line shown in figure 5G, **C)** the unmarked hygR line shown in figure 5I, and **D)** the *Cbunc-119(+)* line shown in Figure 5M. Colored triangles represent *loxP* (yellow), *FRT* (red) and *FRT3* (green) recombinase sites.

Figure S6. Additional TRN, coelomocyte, and unmarked lines

**A-L)** Images of representative L4-staged rRMCE insertion transgenic animals. Shown on top of each set of images is a diagram of the insertion depicting each transcription unit and the plasmid backbone (not to scale). **a)** Widefield view of an L4 animal showing mNG (green) and Scarlet (red) fluorescent signals.

**A-B)** *myo-2p* marked *myo-3p* lines on Chr V. **b-d)** High magnification view of the pharynx of another representative individual: (b) a merge, (c) the Scarlet signal, (d) the mNG signal.

**C-D)** TRN marked *myo-3p* lines on Chr V. **b-c)** high magnification view of the tail of another representative animal: (b) the mNG signal and (c) a merge. **d-e)** High magnification view of the pharynx of another representative individual: (d) a merge and (e) the Scarlet signal.

**E)** TRN marked *nhx-2p* line on chr V. **b-c)** High magnification view of the tail of another representative animal: (b) the mNG signal and (c) a merge. **d-e)** High magnification view of the pharynx of another representative individual: (d) a merge and (e) a long exposure of the Scarlet signal.

**F)** TRN marked *ehs-1p* line on chr V. **b-d)** High magnification view of the head of another representative animal: (b) a merge, (c) a long exposure of the Scarlet signal and (d) a long exposure of the mNG signal.

**G-H)** Coelomocyte marked *mec-4p* mNG and *mex-5p* mNG lines on Chr II. **b-d)** high magnification images of coelomocytes in the mid-body of a representative L4 animals: (b) Scarlet signal, (c) a merge, and (d) mNG signal. **e-f)** High magnification view of the pharynx of another representative individual: (e) a merge and (f) a long exposure of the mNG signal.

**I-J)** Unmarked *rgef-1* promoter mNG lines on Chr IV. **b-d)** High magnification view of the pharynx of another representative individual: (b) a merge and (c) the Scarlet signal, (d) a long exposure of the mNG signal.

**K-L)** Unmarked *myo-3p* mNG lines on Chr IV. **b-d)** High magnification view of the pharynx of another representative individual: (b) a merge and (c) the Scarlet signal, (d) a long exposure of the mNG signal. Note the unexpected GFP(+) cells adjacent to the anterior pharynx in panel K. Strains shown: A) NM6100 *jsSi2107*, B) NM6099 *jsSi2100*, C) NM6103 *jsSi2119*, D) NM6085 *jsSi2110*, E) NM6130 *jsSi2151*, F) NM6125 *jsSi2134*, G) NM6050 *jsSi2055*, H) NM6044 *jsSi2058*, I) NM6144 *jsS2152*, J) NM6139 *jsSi2131*, K) NM6127 *jsSi2142*, L) NM6105 *jsSi2130.* Scale Bars: 100 µm (whole animal images) and 25 µm (pharyngeal image).

Figure S7. Methodology for constructing novel rapid RMCE landing sites

Novel rRMCE landing site can be relatively easily created using either a MosSCI or CRISPR/cas9 insertion strategy {Nonet, 2021, #39572; Nonet, 2020, #20343}. The CRISPR/cas9 strategy was successfully used to create several of the currently available landing sites. However, these specific constructs have not been used to create novel landing sites. To create a novel landing site a targeting plasmid is constructed by Golden Gate assembly using two homology arms (500-1000 bp) adjacent to the targeting site that flank the insertion landing site sequences for either an intermediate landing site (NMp4567 *FRT10 FRT3* landing site) or an rRMCE landing site (NMp4466 *FRT FRT3* landing site). These plasmids are designed to create landing sites that differ slightly from those described in this paper. Specifically, while the landing sites in this paper do not contain a *gpd-2/3 sl2 trans* splice leader *mNG* operonic gene in the landing site, the targeting plasmids created using this strategy will contain this second marker gene. This approach was chosen since a common issue with random insertions in the genome is their incompatibility with germline expression. Lack of germline expression renders the insertion site practically useless. Since the landing sites described in this paper are insertions at sites known to be germline permissive, this was not an issue. However, in creation of novel landing sites, the issue is very significant. Thus, *sl2 mNG* was incorporated into the constructs so that insertions could easily be monitored for germline expression. Since these sequences are excised from the landing site during transgene creation, the inclusion of these sequences has no effect on the structure of the final insertions. Furthermore, if an intermediate landing site is created with *sl2 mNG,* it can still be removed during construction of the final landing site.
